# Covalent Reprogramming of Kinase Binders to Modulate Protein Homeostasis

**DOI:** 10.1101/2025.11.30.691461

**Authors:** Chen Mozes, Xiaokang Jin, Miguel A. Campos, Chen Zhou, Xiaoyu Zhang

## Abstract

Small molecules that modulate protein abundance through induced proximity have expanded the landscape beyond traditional inhibition. Here, we explore how introducing covalent or latent electrophilic groups into a multi-kinase binder scaffold can reprogram protein homeostasis within the kinase family. Using the broad-spectrum kinase ligand TL13-87 as a template, we synthesized analogs bearing α-chloroacetamide, acrylamide, or terminal amine groups. Quantitative proteomics revealed that while most analogs had minimal global impact, MKI-AA uniquely stabilized the mitotic kinase AURKA, a protein often destabilized by ATP-competitive inhibitors. Mechanistic studies showed that MKI-AA acts post-translationally to suppress AURKA ubiquitination and proteasomal degradation. Proteomic mapping of MKI-AA-induced AURKA interactors revealed changes in protein associations upon treatment, providing mechanistic insights into how MKI-AA influences AURKA stability. Intriguingly, adding a short linker to MKI-AA converted it from a stabilizer into a degrader, highlighting how subtle structural variations can invert functional outcomes. These findings demonstrate that electrophilic ligand design can modulate kinase stability and reveal a previously unrecognized mode of covalent proximity-driven protein stabilization.

## Introduction

The controlled modulation of protein abundance by small molecules has emerged as a powerful paradigm in chemical biology and drug discovery. Traditional inhibitors act by blocking enzymatic activity, yet many proteins remain functionally intractable to occupancy-driven pharmacology^[1]^. Targeted protein degradation (TPD) strategies, including heterobifunctional proteolysis targeting chimeras (PROTACs) and monovalent molecular glues, have demonstrated that small molecules can harness the ubiquitin-proteasome system (UPS) to eliminate proteins of interest through induced proximity to E3 ubiquitin ligases^[2]^. More recently, complementary stabilization approaches such as deubiquitinase-targeting chimeras (DUBTACs) have shown that proximity-driven mechanisms can also rescue or enhance protein levels^[3]^. Together, these advances underscore that small molecules can be engineered not only to inhibit, but also to control the stability of target proteins.

An emerging strategy in this area is to introduce covalent or latent electrophilic groups, such as acrylamides, α-chloroacetamides, vinyl sulfonylpiperazines, or terminal amines convertible to aldehydes, into existing ligands to transform them into proximity-inducing agents. Recent studies have shown that such modifications can recruit E3 ligases including DCAF16^[4]^, DCAF11^[4f]^, and FBXO22^[5]^, which recognize electrophilic handles through covalent engagement of reactive cysteine residues. This strategy has enabled the development of both bifunctional and glue-like degraders that induce selective protein degradation, thereby revealing new opportunities to modulate protein homeostasis through induced proximity.

Building on this principle, we sought to investigate how introducing reactive groups, such as α-chloroacetamide, acrylamide, or terminal amine, into multi-kinase binders influences protein homeostasis within the kinase family. Broad-spectrum ATP-competitive inhibitors such as TL13-87 engage multiple kinases through conserved hinge-binding motifs and often tolerate derivatization at the piperazine ring^[6]^. By introducing electrophilic or terminal amine groups to these scaffolds, we hypothesize that the resulting analogs could promote proximity between specific kinase targets and endogenous effector proteins, such as E3 ligases or deubiquitinases, thereby modulating protein homeostasis. Such molecules provide a platform to study how ligand-driven covalency and induced proximity collectively shape kinase abundance, revealing new mechanisms of kinase regulation and expanding the utility of kinase-targeted probes.

## Results and Discussion

### Proteomic characterization of covalent and latent electrophilic analogs of a multi-kinase binder

We selected TL13-87, a multi-kinase binder derived from the ALK-selective inhibitor TAE684, by removing the 2-methoxy group from the 2-anilino hinge-binding segment^[6b, 7]^ (**Figure 1A**). TL13-87 has previously been used to generate heterobifunctional PROTACs through linker attachment at the piperazine ring. The resulting PROTAC shows >90% inhibition of 193 kinases in a 468-kinase panel^[6b]^, demonstrating that it functions as a broadly active tool compound capable of engaging a large number of kinases. To assess whether introducing covalent or latent electrophilic functionalities could affect kinase abundance, we synthesized three TL13-87 analogs bearing an α-chloroacetamide (MKI-CA), acrylamide (MKI-AA), or terminal amine (MKI-A) at the piperazine ring (**Figure 1A**). Using proteomics-based whole-proteome profiling, we evaluated the impact of these analogs on global protein and kinase abundance in HEK293T cells. For comparison, we also included a CRBN-based PROTAC, SK-3-91^[6a]^ (**Figure 1A**).

**Figure 1.**
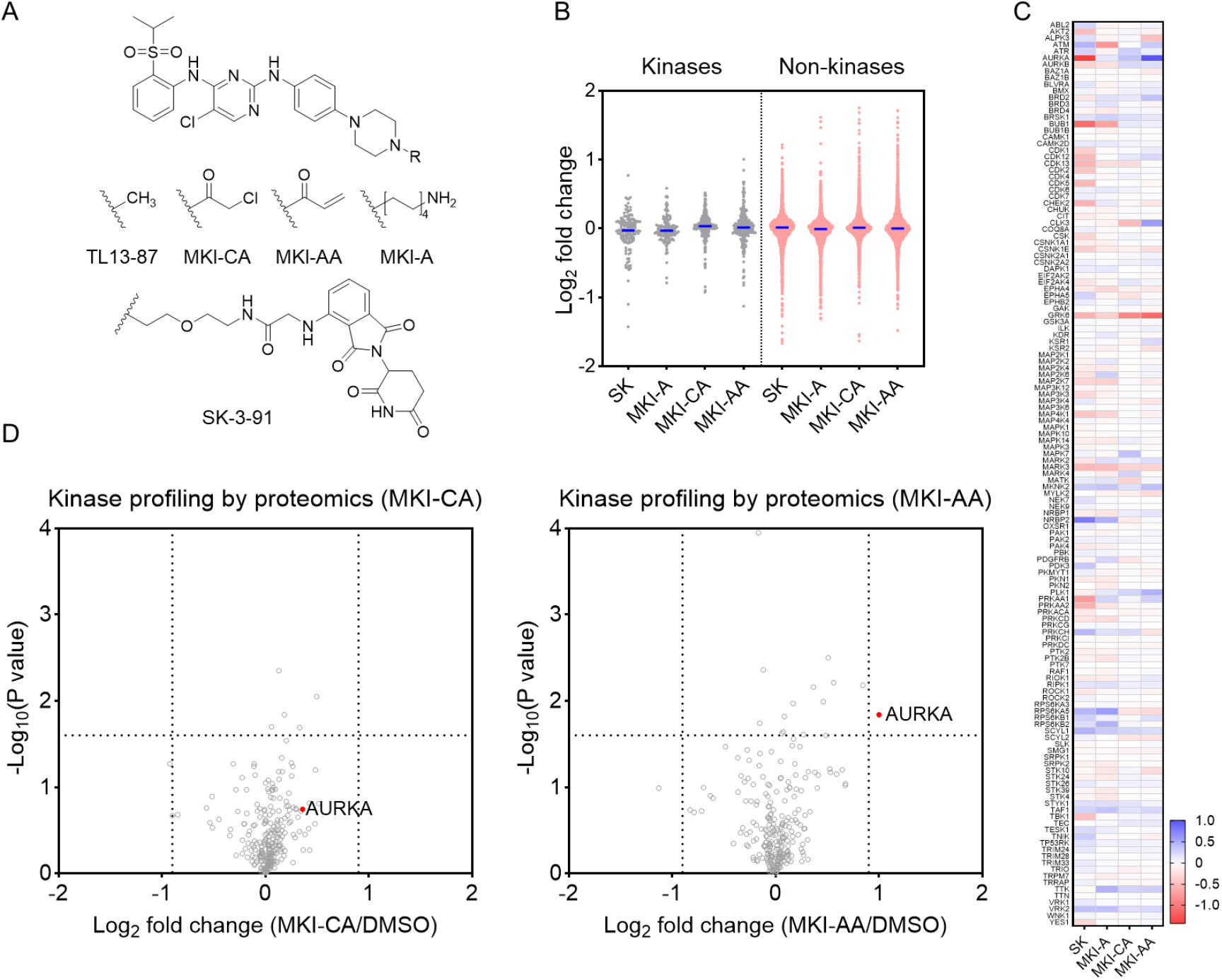
Design and proteomic profiling of multi-kinase-directed probes. **A**. Chemical structures of TL13-87, MKI-CA, MKI-AA, MKI-A, and the CRBN-based PROTAC SK-3-91. **B**. Distribution of protein abundance changes upon probe treatment relative to DMSO control in kinase and non-kinase families. Each dot represents an individual quantified protein. The blue bars indicate the median abundance values. **C**. Heatmap showing the relative abundance changes of quantified kinases following treatment with the indicated kinase-directed probes. **D**. Volcano plots showing global kinome changes upon probe treatment in HEK293T cells (n = 4 biologically independent samples). *P* values were calculated using a two-sided t-test and adjusted for multiple comparisons by the Benjamini-Hochberg method.

Global analysis of quantified kinases and non-kinases showed that none of the compounds caused substantial proteome-or kinome-wide perturbation (**Figure 1B, Figure S1**, and **Table S1**), suggesting that they may not broadly affect protein homeostasis through general effects on translation or the proteasome. Comparative analysis across all quantified kinases revealed that GRK6 abundance was consistently reduced by all four probes, whereas specific kinases such as AURKA and PRKAA1 were selectively down-regulated by SK-3-91 (**Figure 1C** and **Table S1**). Among the TL13-87-derived analogs, MKI-A weakly decreased ATM and BUB1 levels, whereas MKI-CA had minimal impact on the abundance of all quantified kinases except for a mild reduction of GRK6 (**Figure 1C,D**). Notably, MKI-AA selectively stabilized AURKA within the kinase family (**Figure 1C,D**). AURKA is a mitotic serine/threonine kinase critical for spindle assembly and centrosome maturation whose abundance is tightly regulated by proteasomal turnover^[8]^. This observation is intriguing given that AURKA is typically destabilized upon ATP-competitive inhibitor binding^[9]^ or when its stabilizing cofactors, such as NEDD9, are disrupted^[10]^. To our knowledge, there are no prior examples of small molecules that directly stabilize AURKA, suggesting that MKI-AA may engage an atypical binding mode or induce a proximity effect that protects AURKA from turnover.

We next asked whether MKI-AA-mediated AURKA stabilization could tolerate the introduction of an additional linker between the acrylamide and the piperazine ring. To test this, we synthesized two MKI-AA analogs containing either a glycine (MKI-AA2, **Figure 2A**) or a PEG spacer (MKI-AA3, **Figure 2A**) between the acrylamide and the piperazine ring. Notably, incorporation of these short linkers increased cytotoxicity by approximately two-fold in HEK293T cells (**Figure 2B**). Whole-proteome analysis revealed that both MKI-AA2 and MKI-AA3 broadly affected protein abundance, primarily by reducing protein expression levels (**Figure 2C, Figure S2**, and **Table S2**). These results suggested that the modified compounds may have distinct target engagement profiles compared to MKI-AA, leading to substantially altered cellular responses and changes in protein abundance. Interestingly, both analogs lost the ability to stabilize AURKA, and MKI-AA3 exhibited a switch to a potent AURKA destabilizer (**Figure 2D,E**). The AURKA destabilization induced by MKI-AA3 appears to be mediated through a Cullin-RING Ligase E3 ligase and the proteasome system, as co-treatment with the proteasome inhibitor bortezomib (BTZ) or the neddylation inhibitor MLN4924 effectively blocked MKI-AA3-induced AURKA reduction (**Figure 2F**).

**Figure 2.**
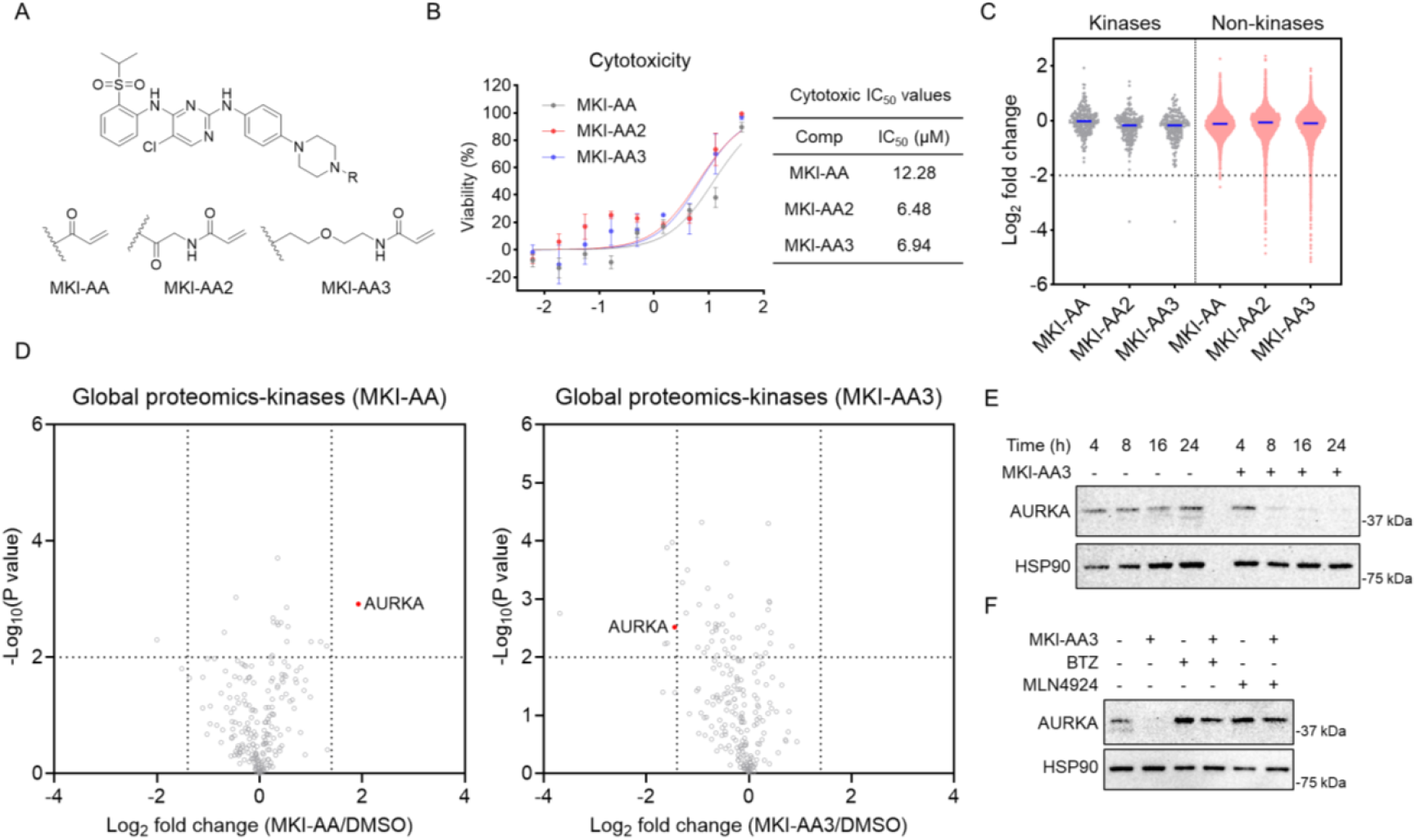
Assessing the impact of linker extension in MKI-AA on AURKA stabilization. **A**. Chemical structures of MKI-AA, MKI-AA2, and MKI-AA3. **B**. Dose-response curves and IC_50_ values of MKI-AA, MKI-AA2 and MKI-AA3 in HEK293T cells. Data are presented as mean ± SEM (n = 3). **C**. Distribution of protein abundance changes upon probe treatment relative to DMSO control in kinase and non-kinase families. **D**. Volcano plots showing global kinome changes upon MKI-AA or MKI-AA3 treatment in HEK293T cells (n = 3 biologically independent samples for MKI-AA3; n = 2 biologically independent samples for MKI-AA). *P* values were calculated using a two-sided t-test and adjusted for multiple comparisons by the Benjamini-Hochberg method. **E**. Western blot analysis of AURKA expression in HEK293T cells following treatment with DMSO or MKI-AA3 (1 µM) for 4-24 hours. The result is representative of two experiments (n = 2 biologically independent samples). **F**. Western blot analysis of AURKA expression in HEK293T cells treated with DMSO or MKI-AA3 (1 µM, 8 hours), with or without co-treatment with BTZ (0.1 µM) or MLN4924 (1 µM). The result is representative of two experiments (n = 2 biologically independent samples).

Recent studies have demonstrated that DCAF11, DCAF16, and FBXO22 are “frequent hitter” E3 ligases that can be engaged by minimal covalent warheads added to protein inhibitors^[4a, 4d, 4f, 4g, 5, 11]^. Given this, we sought to determine whether any of these three ligases are required for MKI-AA-mediated AURKA degradation. We evaluated MKI-AA activity in 22Rv1 *DCAF11* WT and knockout (KO) cells^[12]^ (**Figure S3A**), MDA-MB-231 *DCAF16* WT and KO cells^[13]^ (**Figure S3B**), 22Rv1 *DCAF16* WT and KO cells^[13a]^ (**Figure S3C**), and A549 *FBXO22* WT and KO cells^[14]^ (**Figure S3D**). In all cases, MKI-AA effectively degraded AURKA in the knockout lines. Although it remains possible that dependencies on these E3 ligases may emerge in additional cell contexts not examined here, no such requirement was observed in the cell lines tested in this study. Identifying the essential E3 ligases underlying this mechanism will be an important direction for future work.

### MKI-AA stabilizes AURKA through a post-translational mechanism

To further investigate the mechanism underlying AURKA stabilization, we first validated the effect on endogenous AURKA by Western blot analysis. MKI-AA induced a dose-dependent stabilization of AURKA in HEK293T cells, whereas the α-chloroacetamide analog MKI-CA had no effect (**Figure 3A**). When AURKA was stably overexpressed with GFP or FLAG tags, MKI-AA similarly increased its stability (**Figure 3B,C**), supporting a potential mechanism in which MKI-AA acts directly on AURKA, rather than through secondary pathways that may not apply to overexpressed forms of the protein. To test whether covalency is essential for this effect, we synthesized a non-covalent analog, MKI-PA, which failed to stabilize AURKA (**Figure 3D**), indicating that the covalent acrylamide warhead is required for the observed stabilization. To determine whether this effect occurs post-translationally, we treated cells with cycloheximide (CHX) to block protein synthesis and monitored AURKA degradation over time. Co-treatment with MKI-AA markedly suppressed AURKA degradation (**Figure 3E**), suggesting that MKI-AA likely acts through a post-translational stabilization mechanism. Consistent with this, we observed decreased poly-ubiquitination of FLAG-tagged AURKA upon MKI-AA treatment (**Figure 3F**).

**Figure 3.**
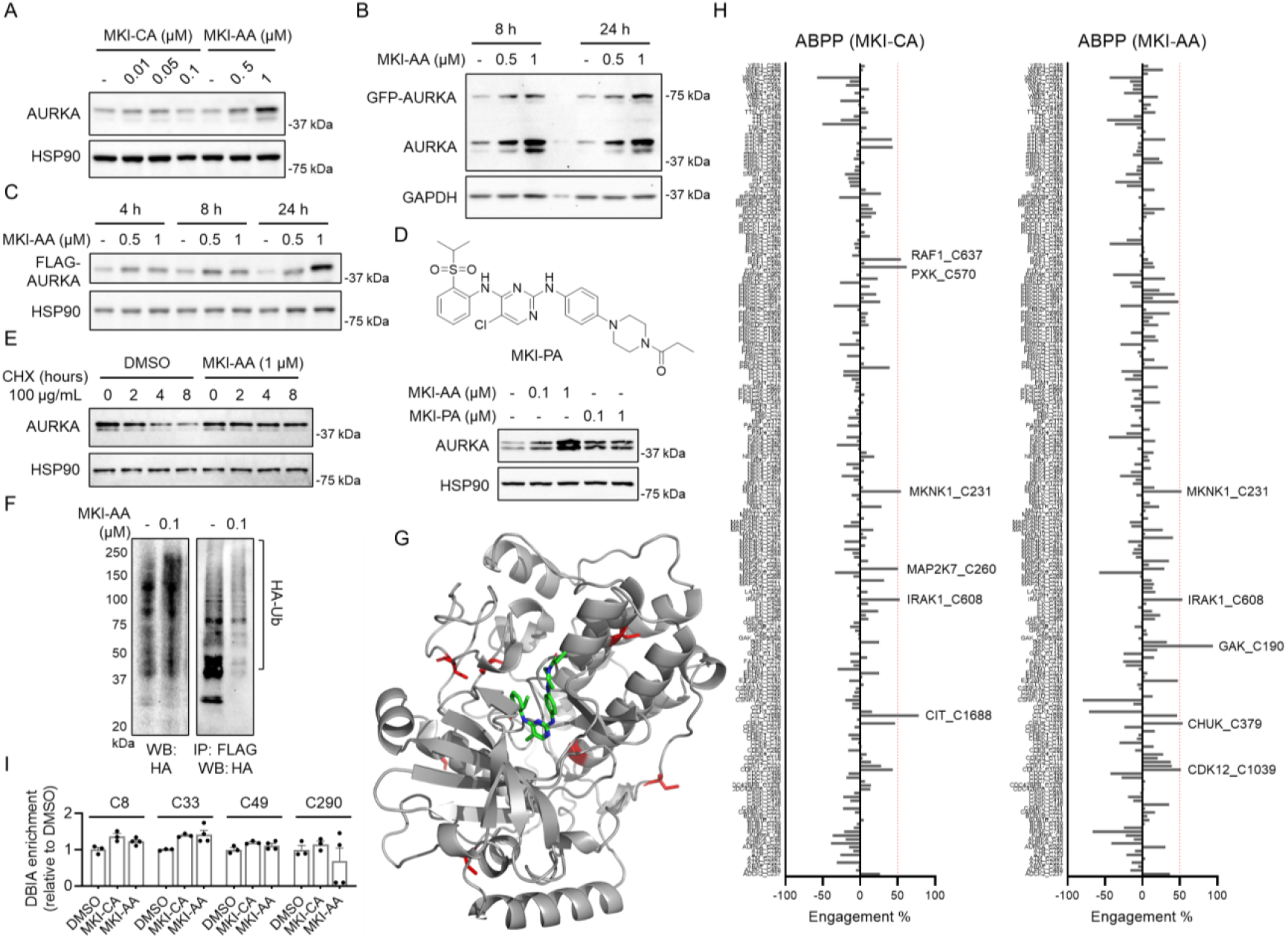
MKI-AA stabilizes AURKA through a post-translational mechanism. **A**. Western blot analysis of AURKA expression in HEK293T cells following treatment with DMSO, MKI-CA or MKI-AA for 8 hours. The result is representative of two experiments (n = 2 biologically independent samples). **B**. Western blot analysis of AURKA expression in HEK293T cells stably expressing GFP-AURKA following treatment with DMSO or MKI-AA for 8 or 24 hours. The result is representative of two experiments (n = 2 biologically independent samples). **C**. Western blot analysis of FLAG-AURKA expression in HEK293T cells stably expressing FLAG-AURKA following treatment with DMSO or MKI-AA for 4, 8 or 24 hours. The result is representative of two experiments (n = 2 biologically independent samples). **D**. Top panel: chemical structure of MKI-PA. Bottom panel: Western blot analysis of AURKA expression in HEK293T cells following treatment with DMSO, MKI-PA or MKI-AA for 8 hours. The result is representative of two experiments (n = 2 biologically independent samples). **E**. Western blot analysis of AURKA expression in HEK293T cells treated with CHX for 2, 4, or 8 hours, with or without co-treatment with MKI-AA (1 µM).The result is representative of two experiments (n = 2 biologically independent samples). **F**. Western blot analysis of AURKA poly-ubiquitination in HEK293T cells expressing HA-ubiquitin and FLAG-AURKA, with or without MKI-AA treatment, followed by FLAG immunoprecipitation and HA blotting. The result is representative of two experiments (n = 2 biologically independent samples). **G**. Structural model generated using the Chai-1 showing the predicted binding pose of MKI-AA (green) within the ATP-binding pocket of AURKA (gray ribbon). The seven cysteine residues in AURKA are highlighted in red. **H**. Target engagement (%) of quantified kinases by MKI-CA or MKI-AA in ABPP experiments. Data are presented as mean (n = 3). **I**. Relative abundance of desthiobiotin iodoacetamide (DBIA)-enriched AURKA peptides with or without treatment with MKI-CA or MKI-AA. Data are presented as mean ± SEM (n = 3).

Mechanistically, MKI-AA likely binds directly to AURKA, as analogs of this scaffold show strong AURKA binding in kinome-wide profiling in a previous study^[6b]^. Molecular modeling performed using the Chai-1 molecular modeling platform^[15]^ supports its fit within the canonical ATP-binding pocket of AURKA, where the acrylamide moiety extends toward the solvent-exposed region (**Figure 3G**). An important question is how the acrylamide group contributes to this effect, specifically, whether it forms a covalent bond with a cysteine residue on AURKA, thereby serving as an intramolecular glue that modulates AURKA’s interactions with other proteins and influences its turnover. However, modeling indicates that all seven cysteine residues of AURKA are distant from the MKI-AA binding pocket (**Figure 3G**), suggesting that direct covalent modification is less likely. This conclusion is further supported by activity-based protein profiling (ABPP), which revealed engagement of multiple cysteines within the kinase family by MKI-AA or MKI-CA (**Figure 3H, Figure S4**, and **Table S3**), but no detectable engagement of the four quantified cysteines in AURKA (**Figure 3I**). Collectively, these results suggest that MKI-AA-induced AURKA stabilization likely involves the covalent engagement of another protein rather than direct modification of AURKA itself.

### Proteomic profiling of MKI-AA-mediated AURKA interactome

Next, we used affinity purification-mass spectrometry (AP-MS) to identify AURKA-interacting partners induced by MKI-AA, aiming to gain insight into potential effector proteins. We employed two strategies. In the first approach, we performed AP-MS under native conditions that preserve protein-protein interactions, enriching FLAG-tagged AURKA from HEK293T cells to compare its interactome with or without MKI-AA treatment. In the second approach, we employed miniTurbo-based proximity labeling^[16]^, which allows the capture of transient or weak interactions that may be missed under native conditions. In this method, AURKA was fused to a miniTurbo tag, and biotin was added to cell culture to label proteins in close proximity to AURKA. In both approaches, SH3GL1 emerged as the top MKI-AA-induced interactor of AURKA (**Figure 4A, Figure S5**, and **Table S4**). Co-immunoprecipitation further validated this finding, as FLAG-AURKA enriched SH3GL1 only in the presence of MKI-AA (**Figure 4B**). To examine whether this interaction involves covalent engagement, we revisited the ABPP dataset (**Table S3**) and found that none of the three quantified cysteines in SH3GL1 (C96, C277, C311) showed significant engagement by MKI-AA (**Figure 4C**). Nevertheless, to test whether SH3GL1 is required for AURKA stabilization, we generated *SH3GL1* KO HEK293T cells and compared AURKA abundance in WT and KO cells upon MKI-AA treatment using both global proteomics and Western blot analysis. However, MKI-AA still stabilized AURKA to a similar extent in *SH3GL1* WT cells (**Figure 4D,E** and **Table S5**).

**Figure 4.**
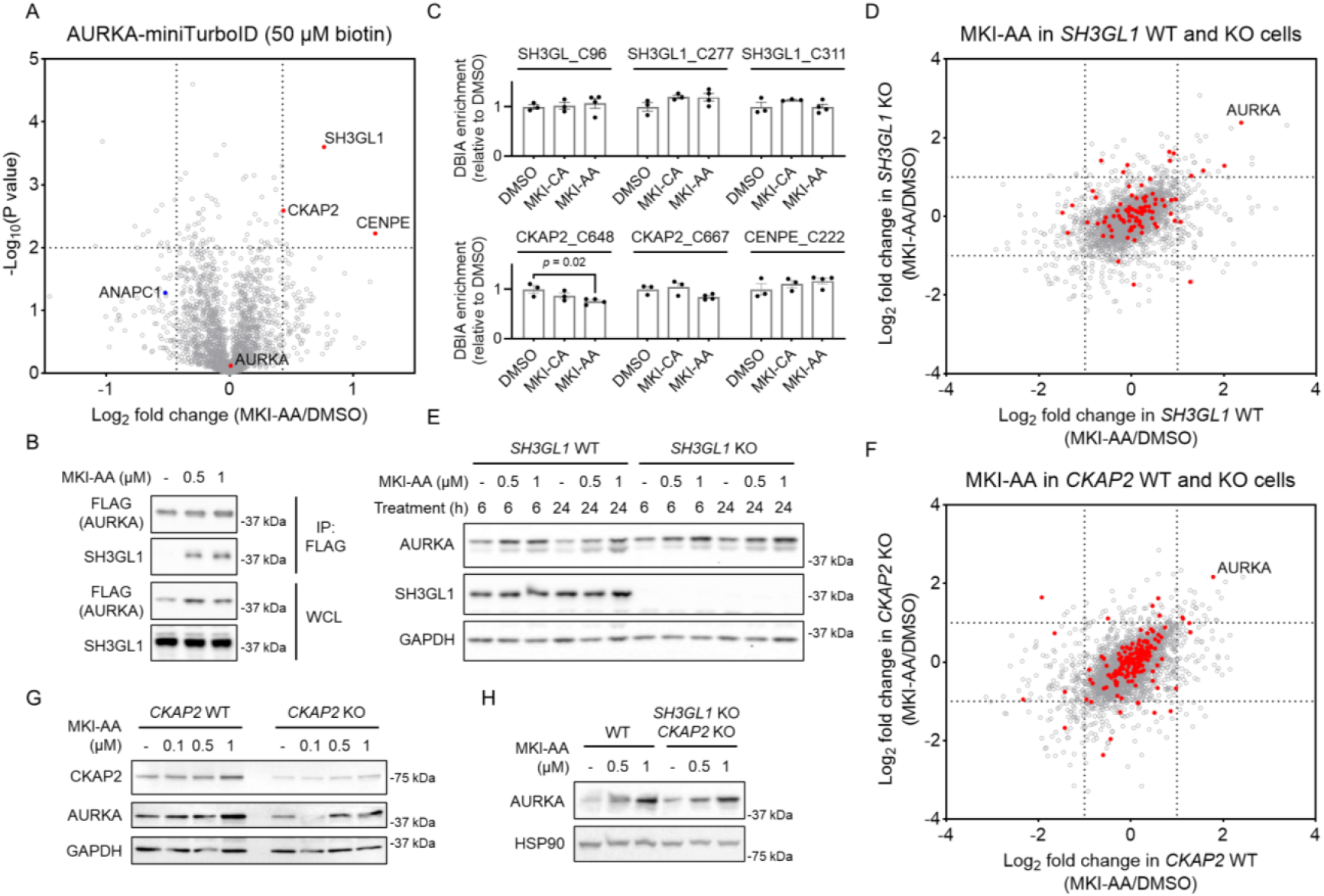
Proteomic profiling of MKI-AA-mediated AURKA interactome. **A**. Volcano plots showing enriched biotinylated proteins comparing MKI-AA versus DMSO treatment in HEK293T cells expressing AURKA-miniTurbo (n = 2 biologically independent samples). *P* values were calculated using a two-sided t-test and adjusted for multiple comparisons by the Benjamini-Hochberg method. **B**. Co-immunoprecipitation analysis showing the interaction between FLAG-tagged AURKA and SH3GL1 with or without treatment with MKI-AA for 2 hours. The result is representative of two independent experiments (n = 2 biologically independent samples). **C**. Relative abundance of DBIA-enriched SH3GL1 and CKAP2 peptides with or without treatment with MKI-CA or MKI-AA. Data are presented as mean ± SEM (n = 3). **D**. Global proteomic comparison of two calculated ratios, MKI-AA/DMSO in *SH3GL1* WT cells and MKI-AA/DMSO in *SH3GL1* KO cells, to identify proteins differentially affected by SH3GL1. Data are presented as mean (n = 3). **E**. Western blot analysis of AURKA expression in HEK293T WT and *SH3GL1* KO cells following treatment with DMSO or MKI-AA for 6 or 24 hours. The result is representative of two experiments (n = 2 biologically independent samples). **F**. Global proteomic comparison of two calculated ratios, MKI-AA/DMSO in *CKAP2* WT cells and MKI-AA/DMSO in *CKAP2* KO cells, to identify proteins differentially affected by CKAP2. Data are presented as mean (n = 3). **G**. Western blot analysis of AURKA expression in HEK293T WT and *CKAP2* KO cells following treatment with DMSO or MKI-AA for 6 hours. The result is representative of two experiments (n = 2 biologically independent samples). **H**. Western blot analysis of AURKA expression in HEK293T WT and *SH3GL1*/*CKAP2* double KO cells following treatment with DMSO or MKI-AA for 6 hours. The result is representative of two experiments (n = 2 biologically independent samples).

We next examined other potential interactors identified by proteomics. The miniTurbo-based proximity labeling approach revealed two additional MKI-AA-induced neo-interactors, CKAP2 and CENPE, that were not identified in the native AP-MS dataset (**Figure 4A** and **Tables S4**). Because *CENPE* is annotated as a common essential gene in the DepMap database^[17]^, generating its knockout was challenging. We therefore focused on CKAP2 and generated *CKAP2* KO HEK293T cells. However, both global proteomics and Western blot analyses showed that MKI-AA still stabilized AURKA in the *CKAP2* KO cells (**Figure 4F,G** and **Table S6**). Next, we generated *SH3GL1*/*CKAP2* double KO cells (**Figure S6**), and MKI-AA-induced AURKA stabilization persisted in the double KO cells (**Figure 4H**). AURKA is known to be targeted for ubiquitin-mediated degradation by the anaphase-promoting complex/cyclosome (APC/C), its native E3 ligase^[18]^. To explore whether MKI-AA might influence this pathway, we re-examined the miniTurbo dataset and found that one APC/C component, ANAPC1, appeared slightly disrupted upon MKI-AA treatment (**Figure 4A**). This finding suggests that MKI-AA may interfere with APC/C-mediated recognition of AURKA, leading to reduced turnover and increased protein stabilization. However, this remains speculative, and other effector proteins or mechanisms may also contribute.

Although the precise mechanism underlying MKI-AA-induced AURKA stabilization remains to be fully elucidated, our study identifies a unique chemical probe that exerts a distinct stabilizing effect on AURKA. This is an outcome uncommon among kinase-directed probes and indicative of an unexpected way in which small molecules can enhance kinase stability, potentially through interference with E3 ligase recognition or a conformational state less susceptible to ubiquitination. Moreover, by stabilizing AURKA through a defined chemical handle, MKI-AA provides a useful tool for studying AURKA-dependent signaling. Moreover, it is noteworthy that while MKI-AA acts as a stabilizer of AURKA, extending its linker by a single PEG unit converts the molecule into a potent degrader. This striking functional switch highlights the balance between stabilization and degradation that can arise from subtle structural modifications. Understanding how such small chemical changes invert the outcome of target engagement will be an important next step, providing mechanistic insights into the principles underlying molecular glue-like mechanisms and the design of new induced-proximity modulators.

## Conclusions

Our study demonstrates that introducing electrophilic handles into multi-kinase scaffolds can reprogram protein fate, leading to either stabilization or destabilization of specific targets. The acrylamide analog MKI-AA uniquely stabilizes AURKA through a post-translational mechanism likely driven by induced proximity, whereas slight structural extensions convert it into a potent degrader. These results highlight that small chemical modifications can toggle between opposing outcomes, providing new insight into how covalent or proximity-based mechanisms can influence kinase abundance and may inspire future strategies for designing chemical probes that modulate kinase stability.

## Acknowledgements

We gratefully acknowledge the support of the NIH R35GM154945 (X.Z.) and the Falk Medical Research Trust Catalyst Award (X.Z.).

## Data Availability Statement

The mass spectrometry proteomics data have been deposited to the ProteomeXchange Consortium via the PRIDE^[19]^ partner repository with the dataset identifier PXD071390.

