## Supplementary Information for "Covalent Reprogramming of Kinase Binders to Modulate Protein Homeostasis"

**Supplementary Tables**

Supplementary Table 1. Global proteomics comparing protein expression in HEK293T cells treated with DMSO, MKI-CA, MKI-AA, MKI-A, or SK-3-91.

Supplementary Table 2. Global proteomics comparing protein expression in HEK293T cells treated with DMSO, MKI-AA, MKI-AA2, or MKI-AA3.

Supplementary Table 3. Cysteine-directed ABPP in HEK293T cells treated with DMSO, MKI-CA, or MKI-AA.

Supplementary Table 4. AP-MS and miniTurbo-based proximity labeling of the AURKA interactome in HEK293T cells treated with DMSO or MKI-AA.

Supplementary Table 5. Global proteomics comparing protein expression in HEK293T parental and *SH3GL1* KO cells treated with DMSO or MKI-AA.

Supplementary Table 6. Global proteomics comparing protein expression in HEK293T parental and *CKAP2* KO cells treated with DMSO or MKI-AA.

### Supplementary Figures

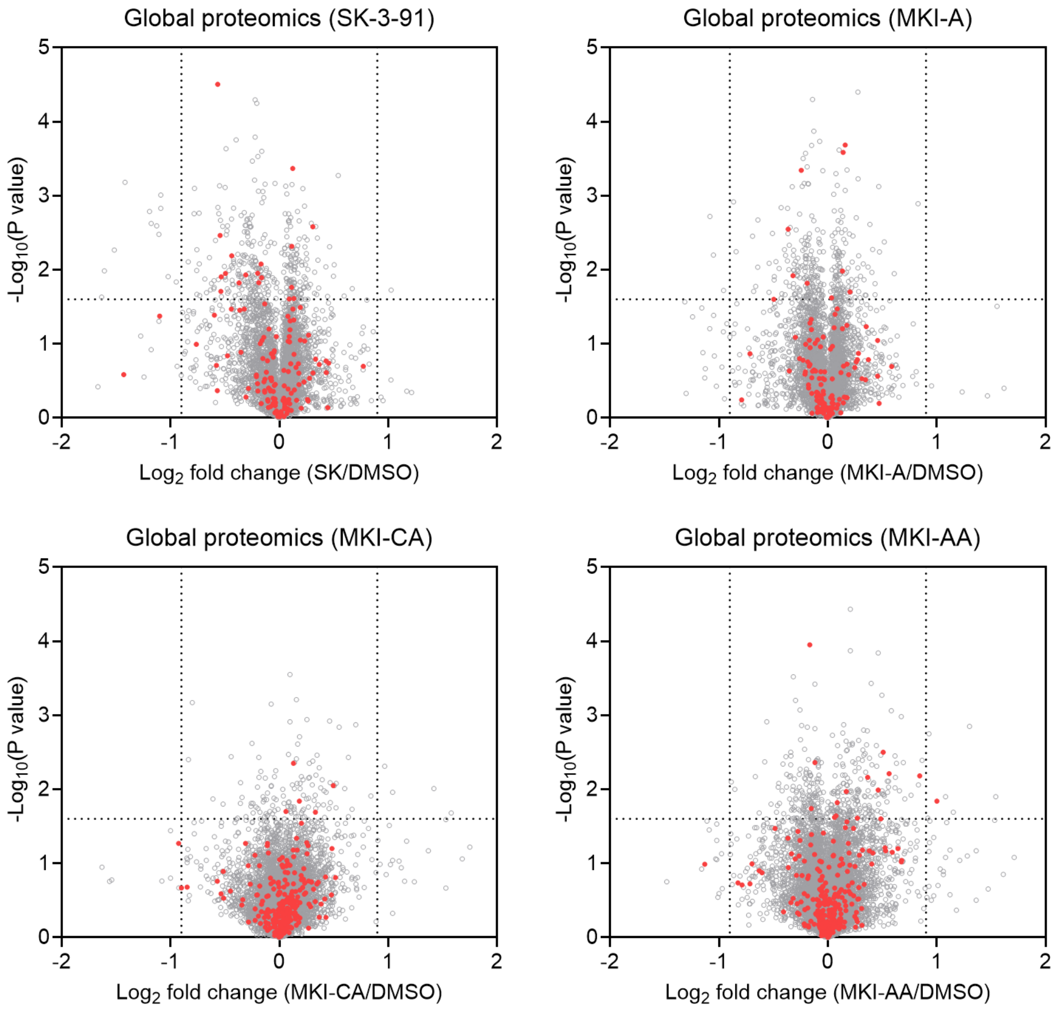


**Figure S1. Proteomic profiling of multi-kinase-directed probes.** Volcano plots showing global proteome changes upon probe treatment in HEK293T cells (n = 4 biologically independent samples for MKI-CA and MKI-AA; n = 2 biologically independent samples for MKI-A and SK-3-91). Kinases are shown in red, and non-kinase proteins in gray. *P* values were calculated using a two-sided t-test and adjusted for multiple comparisons by the Benjamini-Hochberg method.

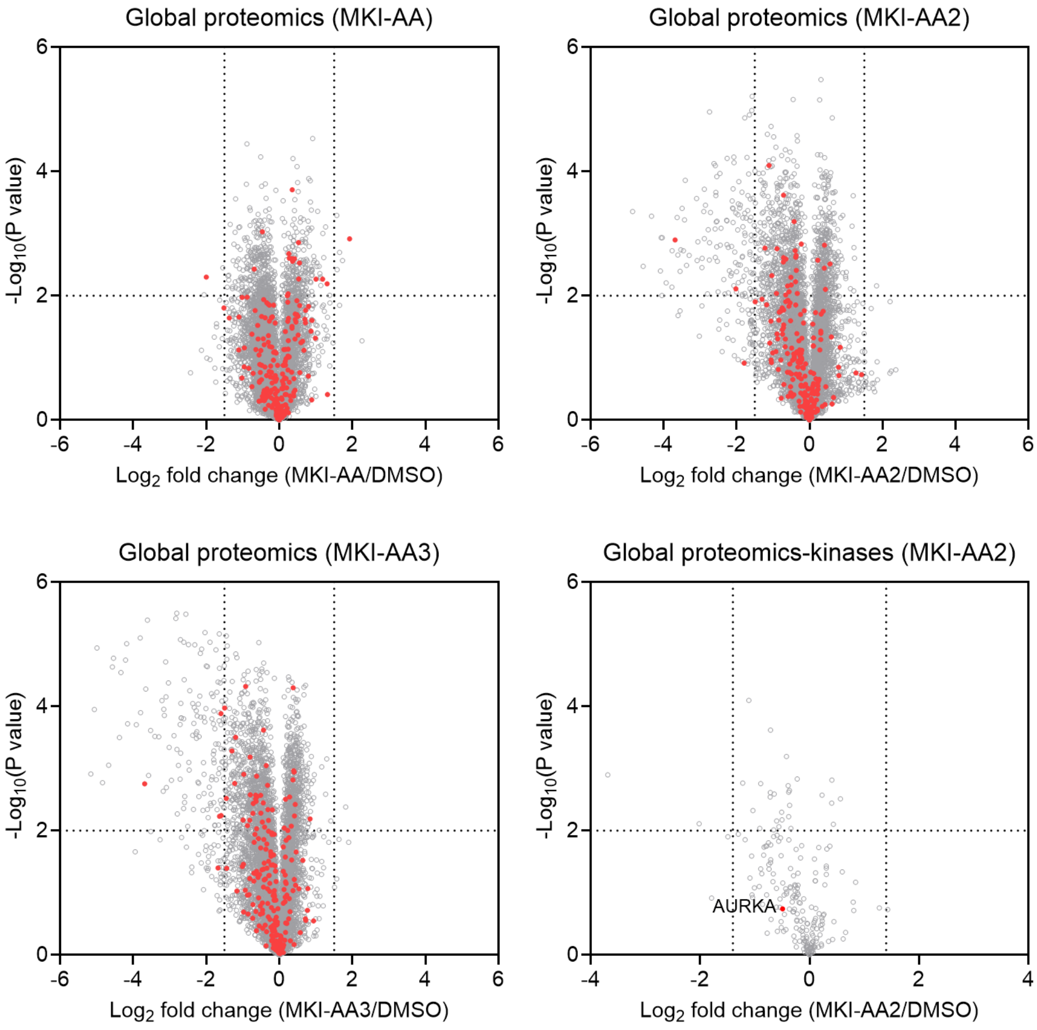


**Figure S2. Proteomic profiling of MKI-AA, MKI-AA2, and MKI-AA3.** Volcano plots showing global proteome (Kinases are shown in red, and non-kinase proteins in gray) or kinome changes upon probe treatment in HEK293T (n = 3 biologically independent samples for MKI-AA3; n = 2 biologically independent samples for MKI-AA and MKI-AA2). *P* values were calculated using a two-sided t-test and adjusted for multiple comparisons by the Benjamini-Hochberg method.

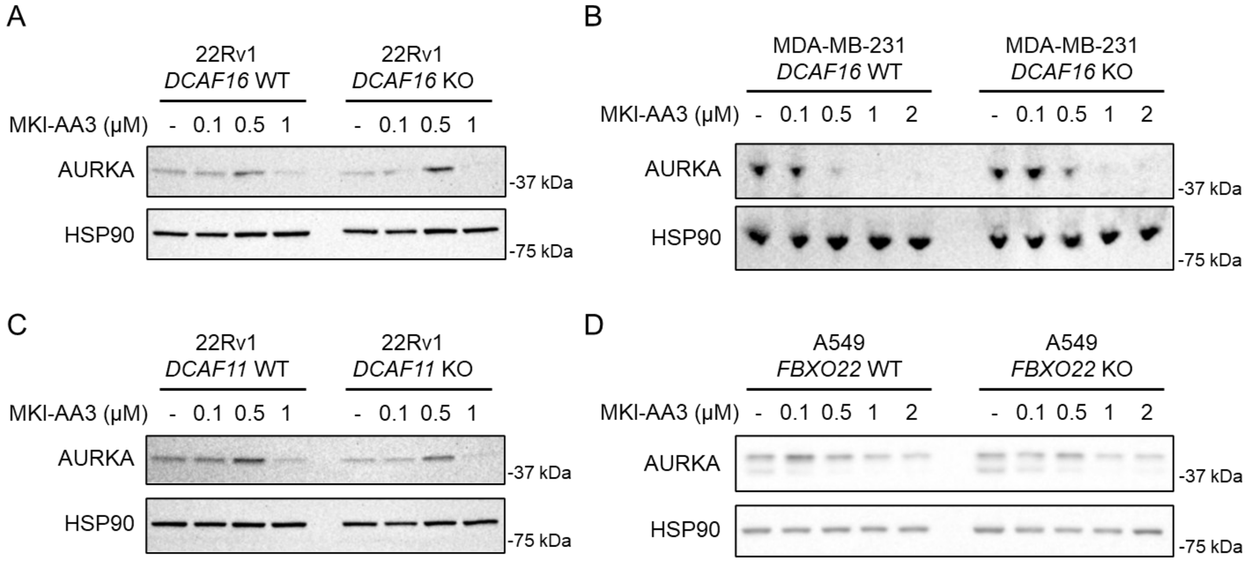


**Figure S3. Assessing the dependence of DCAF11, DCAF16, and FBXO22 on MKI-AA3-mediated AURKA1 degradation. A**. Western blot analysis of AURKA expression in 22Rv1 parental (*DCAF16* WT) and *DCAF16* KO cells following treatment with DMSO or MKI-AA3 for 8 hours. The result is representative of two experiments (n = 2 biologically independent samples). **B**. Western blot analysis of AURKA expression in MDA-MB-231 parental (*DCAF16* WT) and *DCAF16* KO cells following treatment with DMSO or MKI-AA3 for 8 hours. The result is representative of two experiments (n = 2 biologically independent samples). **C**. Western blot analysis of AURKA expression in 22Rv1 parental (*DCAF11* WT) and *DCAF11* KO cells following treatment with DMSO or MKI-AA3 for 8 hours. The result is representative of two experiments (n = 2 biologically independent samples). **D**. Western blot analysis of AURKA expression in A549 parental (*FBXO22* WT) and *FBXO22* KO cells following treatment with DMSO or MKI-AA3 for 8 hours. The result is representative of two experiments (n = 2 biologically independent samples).

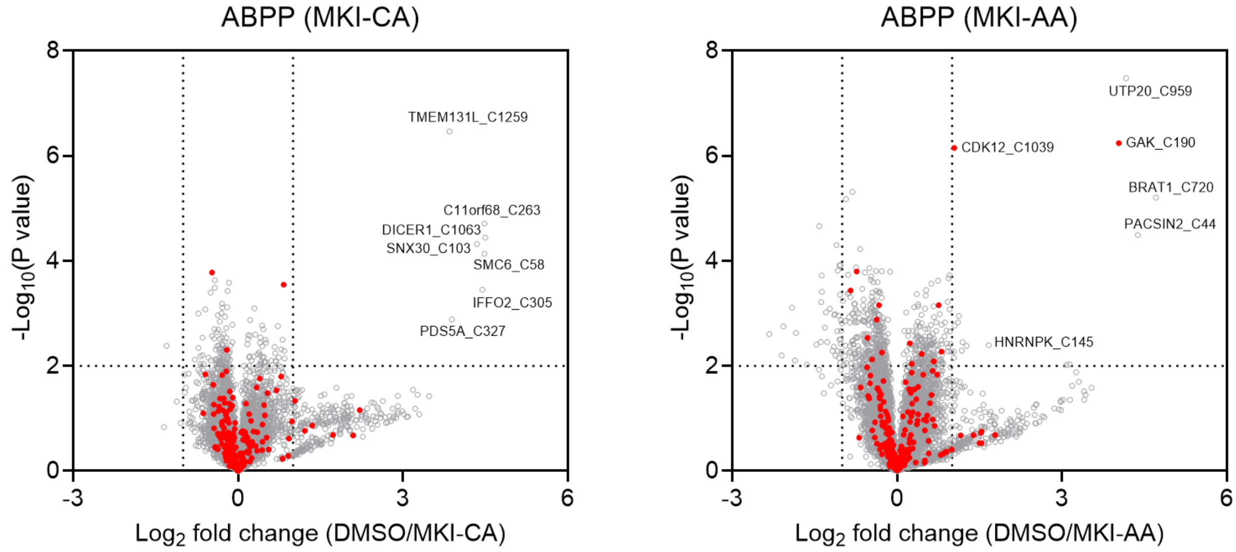


**Figure S4. ABPP analysis of MKI-CA and MKI-AA target engagement.** Volcano plots show global cysteine engagement by MKI-CA or MKI-AA. Kinases are shown in red and non-kinase proteins in gray (n = 3 biologically independent samples). HEK293T cells were treated with MKI-CA (0.2 μM) or MKI-AA (2 μM) for 2 hours. *P* values were calculated using a two-sided t-test and adjusted for multiple comparisons by the Benjamini-Hochberg method.

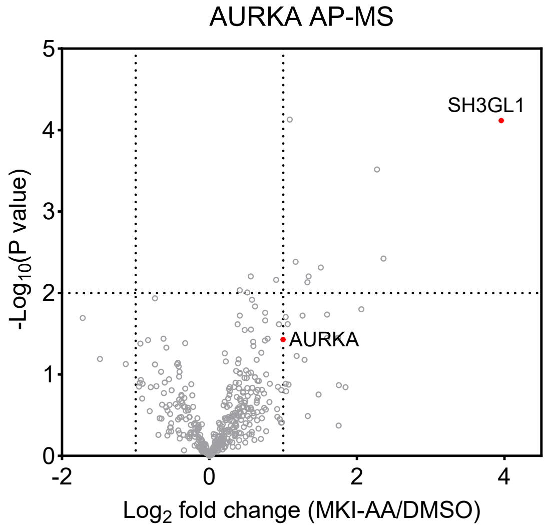


**Figure S5. AP-MS profiling of MKI-AA-mediated AURKA interactome.** Volcano plots showing enrichment of FLAG-AURKA-associated proteins comparing MKI-AA versus DMSO treatment in HEK293T cells (n = 3 biologically independent samples). *P* values were calculated using a two-sided t-test and adjusted for multiple comparisons by the Benjamini-Hochberg method.

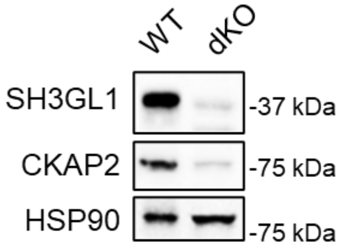


**Figure S6.** Western blot analysis of SH3GL1 and CKAP2 in HEK293T *SH3GL1*/*CKAP2* double KO cells. The result is representative of two experiments (n = 2 biologically independent samples).

### Materials and Methods

#### Reagents

The horseradish peroxidase (HRP)-linked anti-β-actin (clone 13E5, cat# 51255, dilution 1:5000), HRP-linked anti-heat-shock protein 90 (HSP90; clone C45G5, cat# 79641, dilution 1:5000), anti-aurora-A (AURKA; clone D3E4Q, cat# 14475, dilution 1:1000), and HRP-linked rabbit IgG (cat# 7074, dilution 1:1000) antibodies were purchased from Cell Signaling Technologies. The polyclonal CKAP2 antibody (cat# 25486-1-AP, dilution 1:1000) was purchased from Proteintech. The high affinity anti-HA-peroxidase antibody (clone 3F10, cat# 12013819001, dilution 1:5000) was purchased from Millipore Sigma. The anti-FLAG affinity gel (clone M2, cat# A2220), anti-FLAG HRP-conjugated antibody (clone M2, cat# A8592, dilution 1:5000), Biotin (cat# B4639), cycloheximide (cat# C7698) and cOmplete protease inhibitor cocktail (cat#: 11873580001) were purchased from Sigma-Aldrich. Blasticidin (cat# ant-bl-05) and Puromycin (cat# ant-pr-1) were purchased from InvivoGen. Bortezomib (cat# HY-10227) was purchased from MedChemExpress. MLN4924 (cat# 15217) was purchased from Cayman Chemical. Polyethylenimine (molecular weight 40000, cat# 24765-1) was purchased from Polysciences. Enzyme linked chemiluminescence (ECL, cat# 32106) and ECL plus (cat# 32132) western blotting detection reagents, streptavidin agarose (cat# 20349), BCA protein assay kit (cat# 23227), anti-SH3GL1 (polyclonal, cat# 501735533, dilution 1:1000), and tandem mass tag (TMT) isobaric label reagent (cat# 90110) were purchased from Thermo Fisher Scientific. FuGene6 (cat# E2692) transfection reagent and sequencing-grade modified trypsin (cat# V5111) were purchased from Promega. Cas9 endonuclease was purchased from Integrated DNA Technologies (cat# 1081061).

#### Cell Lines

HEK293T, A549, MDA-MB-231, and 22Rv1 cells were obtained from the American Type Culture Collection (ATCC). HEK293T, MDA-MB-231 and A549 were cultured in DMEM (Corning, cat# 15013CV) supplemented with 10% (v/v) Fetal Bovine Serum (Omega

Scientific, cat# FB-01), 2 mM L-glutamine (Gibco, cat# 25030081), and 100 units/mL of a 1:1 mixture of penicillin/streptomycin (Gibco, cat# 15140122). 22Rv1 cells were cultured in RPMI 1640 (Corning) with 10% (v/v) FBS (Omega Scientific) and L-glutamine (2 mM, Gibco). All cell lines tested negative for mycoplasma contamination (ATCC, cat# 30-1012K).

**Cloning, Mutagenesis, and Lentivirus Transduction**

Human AURKA with a N-terminal FLAG tag and human AURKA with a N-terminal FLAG tag followed by miniTurboID were purchased as full-length plasmids from GenScript that cloned in pCDH-CMV-MCS-EF1-puro vector. pHR_dSV40-Aurora A-GFP was obtained from Addgene (plasmid # 67924). Lentivirus was generated by co-transfecting HEK293T cells with a plasmid containing the gene of interest, psPAX2, and pMD2.G using FuGene 6 transfection reagent. The medium containing lentiviral particles was collected after 48 hours, filtered through a 0.45 μm Millex-HV sterile syringe filter unit (MilliporeSigma), and used to transduce HEK293T cells. To generate stable cell lines, puromycin (2 μg/mL) or blasticidin (10 μg/mL) was added and incubated with the cells for 7 days.

**Generation of *SH3GL1* and *CKAP2* Knockout Cells**

HEK293T cells with *SH3GL1* and *CKAP2* CRISPR-Cas9 knockout were generated by electroporating a complex consisting of Cas9 and three sgRNAs using the 4DNucleofector (Lonza Bioscience). The pulse code used was CM-130 for HEK293T cells. Three sgRNAs targeting *SH3GL1* gene (sgRNA#1: UCAACACGGUGUCCAAGAUC; sgRNA#2: GUGCAUGAUCCGCCACGGGA; sgRNA#3: ACCUGCAGCCCAACCCAGGU) were mixed for electroporation. Two sgRNAs targeting *CKAP2* gene (sgRNA#1: GCCAAUGUUACAAUCCGGAA; sgRNA#2: CCACGAUAAGAUCCAAGCAC).

### Generation of Transient Transfection of HA-Ubiquitin in HEK293T Cells Stably Expressing FLAG-AURKA

HEK293T cells stably expressing FLAG-AURKA were grown to 50% confluency in 10 mL DMEM supplemented with 10% fetal bovine serum (FBS), penicillin, streptomycin and glutamine in 10 cm tissue culture dish. 3 µg of DNA was diluted in 1 mL of serum free DMEM and 9 µL of PEI were added. The mixture was incubated at room temperature for 15 min and added dropwise to the cells. Cells were grown for 48 h at 37 °C with 5% $CO_2$ and then were split for treatment followed by FLAG immunoprecipitation and HA blotting.

### Cell Lysis and Western Blot

Cells were lysed in radioimmunoprecipitation lysis buffer (RIPA lysis buffer, Thermo Fisher Scientific, cat# 89900) containing 25 mM Tris-HCl pH 7.6, 150 nM NaCl, 1% sodium deoxycholate, 0.1% SDS and supplemented with cOmplete protease inhibitor cocktail (Millipore Sigma, cat# 11873580001) and Pierce Universal nuclease (Thermo Fisher Scientific, cat# 88701). The suspension mixture was further probe sonicated (40% intensity, 3 rounds, 5 pulses/round). The cell lysates were then centrifuged at 16,000 g at 4 °C for 10 minutes. Protein concentration was determined with a DC assay (BioRad, cat# 5000112). The normalized cell lysates were then mixed in with Laemmli Sample Buffer (BioRad, cat# 1610737EDU) supplemented with 10% β-mercaptoethanol reducing agent (BioRad, cat# 1610710) and heated at 95 °C for 5 minutes. Proteins were separated using 4-20% Novex Tris-Glycine mini gels (Thermo Fisher Scientific, cat# XP04205BOX). The proteins were then transferred onto a 0.2 µM polyvinylidene fluoride (PVDF) membrane (BioRad, cat# 1620177) and incubated with a solution of 5% nonfat milk in TBST buffer (0.1% Tween 20, 20 mM Tris-HCl pH 7.6, and 150 mM NaCl) at room temperature for 1 hour. Antibodies were diluted in 5% nonfat milk in TBST buffer and applied to the membrane at dilutions listed above. After incubation, the membrane was washed 3 times with TBST buffer. Chemiluminescence signals of the membranes were developed using ECL western blotting detection reagent. Signals were captured using ChemiDoc MP (BioRad).

### Cell Viability Assay

Cells were plated in a 96-well clear bottom white plate (Corning) at a density of 100,000 cells per well in 200 µL of DMEM medium treated with varying concentrations of compounds for 24 hours. Following treatment, 50 µL of Cell Titer Glo reagent (Promega) was added to each well and incubated for 10 min at room temperature. Luminescence was measured using CLARIOstar Plus microplate reader (BMG Labtech).

### Immunoprecipitations

Cell pellets were resuspended with NP-40 lysis buffer supplemented with cOmplete protease inhibitor cocktail and Pierce Universal nuclease followed by probe sonication (25% intensity, 2 rounds, 5 pulses/round). The cell suspensions were rotated at 4 °C for 10 minutes before centrifugation at 16,000 g at 4 °C for 10 minutes. Protein concentration was normalized to 2 mg/mL by a DC assay. 30 µL of whole cell lysate samples with normalized concentrations were prepared for western blot analysis by mixing with Laemmli sample buffer and heating at 95 °C for 5 minutes. 500 µL of normalized supernatant solutions were collected for immunoprecipitation. For immunoprecipitation, 30 µL of FLAG affinity gel slurry per sample was added to collected cell lysates and rotated at 4 °C for 2 hours. The affinity gel was then washed 4 times with immunoprecipitation wash buffer (0.2% NP-40, 25 mM Tris-HCl pH 7.4, and 150 mM NaCl). The resulting slurry was then mixed in with Laemmli sample buffer and heated at 95 °C for 5 minutes. The resulting supernatant was collected and subsequently used for western blot analysis.

### Global Proteomics

Cells were lysed in 100 µL of PBS with cOmplete protease inhibitor cocktail using sonication (40% intensity, 3 rounds, 5 pulses/round). Protein concentration was determined with DC assay. A total of 100 µg of protein in 100 µL of lysis buffer was denatured with 8 M urea. For reduction, 5 µL of 200 mM dithiothreitol (DTT) stock solution in water was added, and the mixture was heated to 65 °C for 15 min. Alkylation was performed by adding 5 µL of 400 mM iodoacetamide stock solution in water and incubating in the dark at 37 °C for 30 min. Samples were diluted by adding 300 µL PBS followed by 2 µg of trypsin, and digestion was carried out at 37 °C for 16 h. For TMT labeling, approximately 8.5 µg of each sample in 35 µL of solution was incubated with 9 µL of acetonitrile and 5 µL of TMT tags for 1 hour at room temperature. TMT labeling was quenched by adding 6 µL of a 5% solution of hydroxylamine and incubating for 15 MIN at room temperature. 2.5 µL of formic acid was added to each sample before combining all the samples. Desalting and fractionation were performed using Pierce High pH Reversed-Phase Peptide Fractionation Kit (Thermo Fisher Scientific, Cat# 84868). In brief, the peptide sample was loaded on a spin column, desalted by washing with $H_2O$ with 0.1% formic acid, followed by fractionation with 30 increments of increasing gradient of acetonitrile in 10 mM $NH_4HCO_3$. Every 10th fraction was combined and concentrated resulting in 10 distinct fractions. Peptides were analyzed by liquid chromatography-mass spectrometry (LC-MS) using an Orbitrap Eclipse Tribrid MS coupled with Vanquish Neo UHPLC system. The peptides were introduced to the EASY-Spray HPLC column (C18, 2 µm particle size, 75 µm inner diameter, and 150 mm length) and eluted at a 0.25 µL/min flow rate with the gradient: 5% buffer B (80% $CH_3CN$ with 0.1% FA) in buffer A (water with 0.1% FA) from 0 to 15 minutes, 5% to 45% buffer B from 15-155 minutes, and 45%-100% buffer B from 155-180 minutes. Voltage of the nano-LC electrospray ionization source set to 1.5 kV. The analysis started with an MS1 master scan (Orbitrap analysis; resolution 60,000; m/z range 375-1600; RF lens 30%; standard automatic again control (AGC) target; auto maximum injection time). For MS2 analysis, initial precursor ions were isolated by the quadrupole with an isolation window of 0.7 and then subjected to higher-energy collisional dissociation (HCD) in the ion trap (stand AGC; collision energy 27%; maximum injection time 35 ms). After each MS2 spectrum, synchronous precursor selection (SPS) chose up to 10 MS2 fragment ions for MS3 analysis. These precursors were once again fragmented by HCD and analyzed by the Orbitrap (AGC 250%; collision energy 55%; maximum injection time 200 ms; resolution 60,000). The raw data was collected using Xcalibur (version 4.5.445.18).

Raw mass spectrometry data were analyzed using Proteome Discoverer 2.5 (Thermo Scientific). Spectra were searched with the Sequest HT algorithm against the UniProt human reference proteome (UP000005640_9606_Human.fasta). Searches were performed with fully tryptic cleavage specificity, a precursor mass tolerance of 10 ppm, and a fragment ion tolerance of 0.6 Da. Fixed modifications included carbamidomethylation of cysteines and TMT labeling on peptide N-termini and lysine residues. Methionine oxidation and protein N-terminal acetylation were specified as variable modifications. Peptide-spectrum matches were evaluated using Percolator, and identifications were filtered to a 1% false-discovery rate at both the peptide and protein levels. Reporter ion intensities were extracted from SPS-MS3 scans with a 20 ppm integration window. Quantification was carried out using unique and razor peptides, with total peptide-signal normalization applied across all TMT channels.

**Cysteine-Directed Activity-Based Protein Profiling (ABPP)**

Cells were suspended in 500 µL of PBS and lysed through probe sonification (40% intensity, 3 rounds, 5 pulses/round). The cell lysates were then centrifuged at 16,000 g at 4 °C for 10 minutes. Protein concentration was normalized to 1 mg/mL by a DC assay. 500 µL of normalized cell lysates were labeled with 100 µM desthiobiotin iodoactamide (DBIA) by incubating for 1 hour at room temperature. Protein cleanup was performed by adding 100 µL of a 1:1 mixture of hydrophobic:hydrophilic Sera-Mag SpeedBeads to each sample. The cell lysates were incubated with the beads at room temperature for 5 minutes (1,000 rpm rotation in a thermomixer). The lysate-bead mixture was then incubated after adding 1 mL of absolute ethanol at room temperature for 5 minutes (1,000 rpm rotation in a thermomixer). The supernatant was aspirated after beads had settled using a DynaMag2 magnet. The beads were resuspended in 500 µL of a 2 M solution of urea in PBS. The proteins were then reduced with 25 µL of DTT (200 mM in HPLC-grade water) by incubating at 65 °C for 15 minutes. Alkylation was then performed by adding 25 µL of IA (400 mM in HPLC-grade water) and incubating in the dark at 37 °C for 30 minutes. The beads were washed with 1 mL of absolute ethanol three times, resuspending in 200 µL of PBS and digested with 2 µg of trypsin at 37 °C for 16 hours. After digestion, the supernatant was collected and incubated with 300 µL of ABPP wash buffer (50 mM TEAB, 150 mM NaCl, 0.2% NP-40) containing 50 µL of streptavidin agarose. The streptavidin-peptide mixture was rotated at room temperature for 2 hours. Upon completion, the beads were washed with ABPP wash buffer (1 mL x 3), PBS (1 mL x 3), and with HPLC-grade water (1 mL x 3), in a BioSpin column. Peptides were eluted from the beads by adding 300 µL of 50% acetonitrile with 0.1% formic acid in HPLC-grade water. The eluted peptides were dried with a SpeedVac vacuum concentrator. Samples were resuspended in 100 µL of 100 mM TEAB in 30% dry acetonitrile. For TMT labeling, added to each sample 3 µL of TMT tags and incubated for 1 hour at room temperature. TMT labeling was quenched by adding 3 µL of a 5% solution of hydroxylamine and incubating for 15 minutes at room temperature. 5 µL of formic acid was added to each sample before combining all the samples. Desalting and fractionation were performed using Pierce High pH Reversed-Phase Peptide Fractionation Kit. In brief, the peptide sample was loaded on a spin column, desalted by washing with $H_2O$ with 0.1% formic acid, followed by fractionation with 15 increments of increasing gradient of acetonitrile in 10 mM $NH_4HCO_3$. Every 5th fraction was combined and concentrated resulting in 5 distinct fractions. Peptides were analyzed by LC-MS as described above.

**Affinity Purification Mass Spectrometry (AP-MS)**

Cells were lysed in NP-40 lysis buffer with the cOmplete protease inhibitor cocktail. After centrifugation at 16,000 g for 10 minutes at 4 °C, the supernatant was collected for immunoprecipitation. Protein lysates were incubated with 25 µL FLAG affinity gel slurry per sample for 2 hours at 4 °C. The gel was washed four times in an immunoprecipitation washing buffer, followed by two washes with PBS. FLAG-AURKA and the associated proteins were eluted by heating the beads at 65 °C for 10 minutes in 8 M urea in PBS. The eluted proteins were then reduced with 12.5 mM DTT at 65 °C for 15 minutes, followed by alkylation with 25 mM iodoacetamide at 37 °C for 30 minutes. The protein solution was diluted in PBS to achieve a urea concentration of 2 M and digested with 2 µg of trypsin at 37 °C for 16 hours. 6 µL TMT tags were added and incubated at room temperature for 1 hour, after which the reaction was quenched with 6 μL of a 5% hydroxylamine solution and 2.5 μL of formic acid. Samples were then pooled and desalted using a Sep-Pak C18 cartridge (Waters, cat# WAT054955). The eluted peptide solution was dried with a SpeedVac concentrator and analyzed by LC-MS as described above.

**Proximity Labeling with miniTurboID**

HEK293T cells stably expressing FLAG-AURKA-miniTurboID were treated with DMSO or probe in the presence of 50 μM biotin (Sigma-Aldrich, cat# B4639) for 6 hours before being collected and washed with PBS. Cell pellets were resuspended with PBS and lysed with probe sonication (40% intensity, 3 rounds, 5 pulses/round). Protein concentration was measured by DC assay. A total of 1 mg of protein in 500 μL of lysis buffer was denatured with freshly made 8 M urea in PBS and 10 μL of 10% SDS. For reduction, 25 μL of a 200 mM DTT solution in water was added, and the sample was heated to 65 °C for 15 minutes. Alkylation was performed by adding 25 μL of a 400 mM iodoacetamide solution in water, followed by incubation in the dark at 37 °C for 30 minutes. Added 100 μL of 10% SDS and transferred each sample to a 15 mL tube with 5 mL PBS. For streptavidin enrichment, 50 μL streptavidin beads per sample were washed, added to each sample and rotated for 1.5 hours. The beads were then washed with 0.2% SDS/PBS (1 mL x 2), PBS (1 mL x 2), HPLC water (1 mL x 2) and 100 mM TEAB (1 mL x 1). After removing the supernatant, the beads were resuspended with 70 μL 1 M urea in 100 mM TEAB and digested with 2 μg of trypsin at 37 °C for 16 hours. The supernatant was collected followed by addition of 25 μL acetonitrile and 5 μL of TMT labels. The resulting mixture was allowed to incubate at room temperature for 1 hour. TMT labeling was quenched by adding 6 μL of a 5% solution of hydroxylamine and incubating for 15 minutes at room temperature. 5 μL of formic acid was added to each sample before combining and drying with a SpeedVac vacuum concentrator. The pooled peptides were subjected to desalting using a Sep-Pak C18 cartridge and the resulting elution was dried with a SpeedVac vacuum concentrator. Peptides were analyzed by LC-MS as described above.

**Statistical Analysis**

Quantitative data is presented as scatter plots, with the mean displayed alongside standard error of the mean (SEM) as error bars. Comparisons between two groups were analyzed using an unpaired two-tailed Student's t-test.

**Chemistry**

All reactions were carried out under a nitrogen atmosphere in flamed-dried glassware with magnetic stirring unless stated otherwise. Chemicals and reagents were purchased from a variety of vendors, including Sigma Aldrich, Thermo Fisher Scientific, Ambeed, CombiBlocks, and were used without further purification, unless noted otherwise. Anhydrous solvents were obtained as commercially available pre-dried, oxygen-free formulations. Normal and reverse phase purification of reaction products was carried out by flash chromatography on Biotage Selekt systems with ultragrade silica cartridges. Preparative thin layer chromatography (PTLC) was carried out using glass backed PTLC 20x20 cm plates 250 or 500 μm thickness (Miles Scientific). Analytical thin-layer chromatography was performed on 0.25 mm silica gel 60-F plates. Visualization was done with UV light and/or by ninhydrin or $KMnO_4$ staining. $^1$H-NMR spectra were recorded on a Bruker AVANCE III 500 MHz with DCH Cryoprobe (500 MHz) spectrometer and are reported in ppm using solvent as an internal standard ($CDCl_3$ at 7.26 ppm, CDCl3 with 0.03% v/v TMS at 7.26 ppm, MeOD at 3.31 ppm). Data are reported as (bs = broad singlet, s = singlet, d = doublet, t = triplet, q = quartet, m = multiplet, etc.; coupling constant(s) in Hz; integration) Proton-decoupled $^{13}$C NMR spectra were recorded on Bruker AVANCE III 500 MHz with DCH Cryoprobe (126 MHz) spectrometer and are reported in ppm using residual solvent as an internal standard ($CDCl_3$ at 7.26 ppm, CDCl3 with 0.03% v/v TMS at 7.26 ppm, MeOD at 3.31 ppm). Low-resolution mass spectra (not reported herein) were obtained on a WATERS Acquity I-Class UPLC-MS with 17 a single quad detector (ESI), ELSD, and PDA. High resolution mass spectra were obtained using an Agilent 6201 MSLC-TOF (ESI).

### Synthetic Procedures

The synthesis of 5-chloro-N4-(2-(isopropylsulfonyl)phenyl)-N2-(4-(piperazin-1-yl)phenyl)pyrimidin-e-2,4-diamine (**MKI-NH**) was performed according to a previously reported procedure.[1]

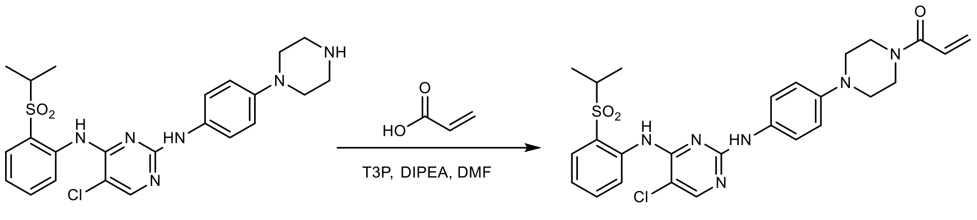


#### 1-(4-(4-((5-chloro-4-((2-(isopropylsulfonyl)phenyl)amino)pyrimidin-2-yl)amino)phenyl)piperazin-1-yl)prop-2-en-1-one (MKI-AA)

A solution of acrylic acid (3.3 μL, 0.048 mmol, 1.2 equiv.) and propanephosphonic acid anhydride (T3P) (43 mg, 0.051 mmol, 1.7 equiv.) in dry DMF (2 mL) under $N_2$ was cooled to 0 °C and stirred for 10 minutes. Then diisopropylethylamine (20.9 μL, 0.12 mmol, 3 equiv.) was added followed by amine MKI-NH (20 mg, 0.04 mmol, 1 equiv.) and the mixture was stirred at 0 °C for another 30 min, then at room temperature for overnight. The reaction mixture was concentrated under reduced pressure and purified by preparative TLC (hexane/ethyl acetate 1:4) to yield **MKI-AA** as a pale yellow solid (10 mg, 18.5 μmol, 46%). $^1$H-NMR (500 MHz, $CDCl_3$): δ 9.63 (s, 1H), 8.58 (d, *J* = 8.5 Hz, 1H), 8.10 (s, 1H), 7.90 (dd, *J* = 8.0, 1.5 Hz, 1H), 7.56 (t, *J* = 7.0 Hz, 1H), 7.42 (d, *J* = 9.0 Hz, 2H), 7.24 (t, *J* = 8.0 Hz, 2H), 6.90 (d, *J* = 8.5 Hz, 2H), 6.61 (dd, *J* = 17.0, 10.5 Hz, 1H), 6.33 (dd, *J* = 17.0, 2.0 Hz, 1H), 5.74 (dd, *J* = 10.5, 2.0 Hz, 1H), 3.87 (bs, 2H), 3.74 (bs, 2H), 3.26-3.20 (m, 1H), 3.15 (t, *J* = 5.5 Hz, 4H), 1.31 (d, *J* = 7.0 Hz, 6H). $^{13}$C-NMR (126 MHz, $CDCl_3$): δ 165.54, 158.06, 155.56, 155.06, 147.31, 138.54, 134.54, 132.60, 131.40, 128.39, 127.41, 124.63, 123.60, 123.27, 122.08, 117.58, 105.99, 55.76, 50.70, 50.17, 45.92, 42.04, 15.49. HRMS (ESI+) m/z calculated for $C_{26}H_{29}ClN_6O_3S$ $[M+H]^+$: 541.17831; found: 541.17658.

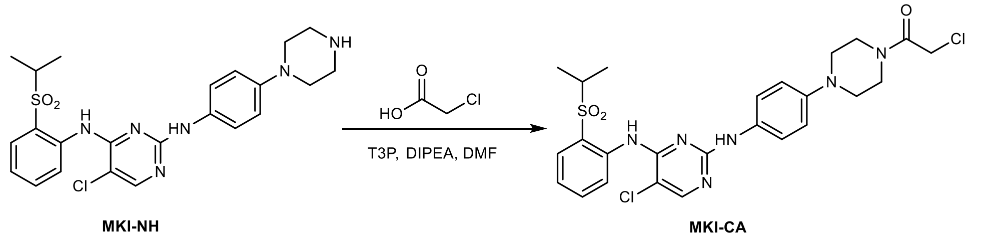


**2-chloro-1-(4-(4-((5-chloro-4-((2-(isopropylsulfonyl)phenyl)amino)pyrimidin-2-yl)amino)phenyl)piperazin-1-yl)ethenone (MKI-CA)**

This compound was synthesized with the same procedure as MKI-AA, starting from chloroacetic acid (2.9 µL, 0.048 mmol, 1.2 equiv.). The preparative TLC was run with hexane/ethyl acetate 1:4 to yield **MKI-CA** as a yellow solid (7.1 mg, 12.6 µmol, 31%). $^{1}$H-NMR (500 MHz, $CDCl_3$): δ 9.95 (s, 1H), 8.58 (d, *J* = 8.5 Hz, 1H), 8.11 (s, 1H), 7.90 (dd, *J* = 8.0, 1.5 Hz, 1H), 7.57 (t, *J* = 7.0 Hz, 1H), 7.42 (d, *J* = 9.0 Hz, 2H), 7.25 (t, *J* = 8.5 Hz, 2H), 7.10 (bs, 1H), 6.91 (d, *J* = 9.0 Hz, 2H), 4.12 (s, 2H), 3.81 (m, 2H), 3.71 (m, 2H), 3.28-3.22 (m, 1H), 3.20 (m, 2H), 3.15 (m, 2H), 1.31 (d, *J* = 7.0 Hz, 6H). $^{13}$C-NMR (126 MHz, $CDCl_3$): δ 165.28, 157.89, 155.58, 154.83, 147.20, 138.51, 134.55, 132.67, 131.44, 124.68, 123.57, 123.34, 122.08, 117.73, 106.17, 55.79, 50.51, 50.09, 46.44, 42.27, 40.95, 15.50. HRMS (ESI+) m/z calculated for $C_{25}H_{28}Cl_2N_6O_3S$ $[M+H]^+$: 563.13934; found: 563.14062.

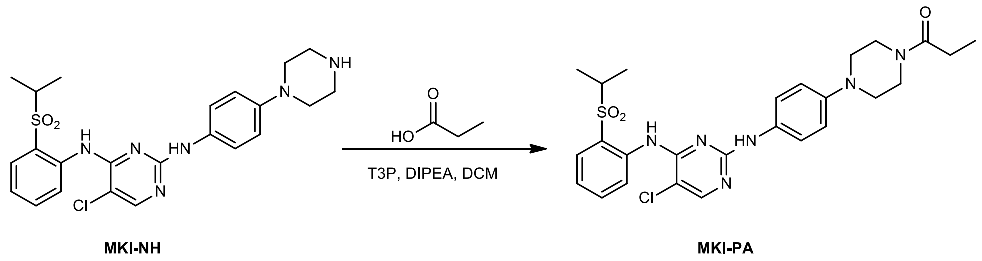


**1-(4-(4-((5-chloro-4-((2-(isopropylsulfonyl)phenyl)amino)pyrimidin-2-yl)amino)phenyl)piperazin-1-yl)propan-1-one (MKI-PA)**

This compound was synthesized with the same procedure as MKI-AA, starting from propionic acid (3.8 µL, 0.048 mmol, 1.2 equiv.). The preparative TLC was run with hexane/ethyl acetate 1:4 to yield **MKI-PA** as a brown solid (13 mg, 24 µmol, 60%). $^1$H-NMR (500 MHz, $CDCl_3$): 9.63 (s, 1H), 8.58 (d, *J* = 8.5 Hz, 1H), 8.09 (s, 1H), 7.90 (dd, *J* = 8.0, 1.5 Hz, 1H), 7.55 (dt, *J* = 8.0, 1.5 Hz, 1H), 7.42 (d, *J* = 9.0 Hz, 2H), 7.35 (bs, 1H), 7.24 (dt, *J* = 7.5, 1.0 Hz, 1H), 6.90 (d, *J* = 9.0 Hz, 2H), 3.81-3.78 (m, 2H), 3.65-3.62 (m, 2H), 3.27-3.18 (m, 1H), 3.14-3.09 (m, 4H), 2.40 (q, *J* = 7.5 Hz, 2H), 1.31 (d, *J* = 7.0 Hz, 6H), 1.18 (t, *J* = 7.5 Hz, 3H). $^{13}$C-NMR (126 MHz, $CDCl_3$): δ 172.49, 158.03, 155.57, 154.93, 147.40, 138.52, 134.54, 132.53, 131.39, 124.63, 123.62, 123.28, 122.09, 117.53, 105.93, 55.76, 50.60, 50.24, 45.52, 41.65, 26.62, 15.48, 9.62. HRMS (ESI+) m/z calculated for $C_{26}H_{31}ClN_6O_3S$ $[M+H]^+$: 543.19397; found: 543.19356

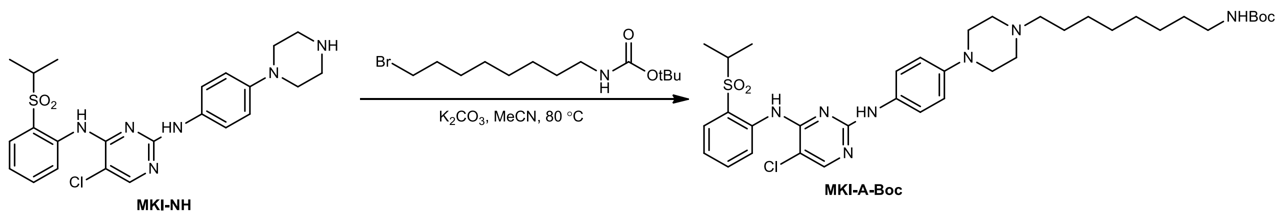


**tert-butyl-(8-(4-(4-((5-chloro-4-((2-(isopropylsulfonyl)phenyl)amino)pyrimidin-2-yl)amino)phenyl)piperazin-1-yl)octyl)carbamate (MKI-A-Boc)**

MKI-NH (100 mg, 0.17 mmol, 1 equiv.), N-Boc-8-bromooctane-amine (63 mg, 0.2 mmol, 1.2 equiv.) and potassium carbonate (71 mg, 0.5 mmol, 3 equiv.) were dissolved with 2.5 acetonitrile and refluxed at 80 °C for overnight. The mixture was diluted with dichloromethane and filtered. The filtrate was concentrated under reduced pressure and purified by flash chromatography (10%-100% EtOAc in hexane) to yield **MKI-A-Boc** as an off-white solid (54 mg, 76 µmol, 44%). $^1$H-NMR (500 MHz, $CDCl_3$): δ 9.61 (s, 1H), 8.59 (d, *J* = 8.0 Hz, 1H), 8.10 (s, 1H), 7.88 (d, *J* = 7.5 Hz, 1H), 7.54 (t, *J* = 8.0 Hz, 1H), 7.37 (d, *J* = 7.0 Hz, 2H), 7.22 (t, *J* = 7.5 Hz, 1H), 6.92-6.89 (m, 3H), 4.50 (bs, 1H), 3.26-3.21 (m, 1H), 3.20-3.14 (m, 4H), 3.13-3.08 (m, 2H), 2.63-2.60 (m 4H), 2.41-2.37 (m, 2H), 1.55-1.50 (m, 2H), 1.44 (s, 9H), 1.31 (s, 10H), 1.30 (s, 6H). $^{13}$C-NMR (126 MHz, $CDCl_3$): δ 158.23, 155.99, 155.34, 155.30, 147.92, 138.53, 134.48, 131.44, 131.20, 124.30, 123.41,

122.96, 122.27, 116.66, 105.75, 79.03, 58.83, 55.58, 53.33, 49.79, 40.63, 30.08, 29.50, 29.23, 28.45, 27.54, 26.92, 26.76, 15.37.

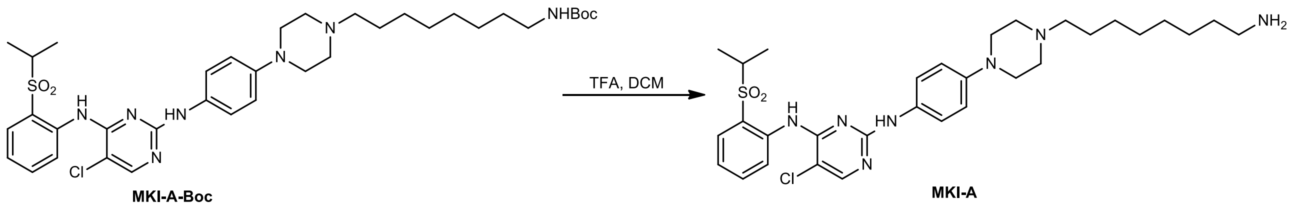

MKI-A-Boc MKI-A

**N2-(4-(4-(8-aminooctyl)piperazin-1-yl)phenyl)-5-chloro-N4-(2-(isopropylsulfonyl)phenyl)pyrimidine-2,4-diamine (MKI-A)**

MKI-A-Boc (54 mg, 0.076 mmol, 1 equiv.) was dissolved with DCM (240 µL) followed by a dropwise addition of TFA (60 µL). The mixture was stirred overnight at room temperature and then concentrated under reduced pressure. The crude mixture was purified with reverse-phase chromatography on a Biotage C18 column (acetonitrile in $H_2O$, 5-95% gradient) to afford **MKI-A** as a yellow solid (10 mg, 16 µmol, 21%). $^1$H-NMR (500 MHz, MeOD): δ 8.54-8.45 (m, 1H), 8.14 (s, 1H), 7.92 (d, *J* = 8.0 Hz, 1H), 7.68 (t, *J* = 7.5 Hz, 1H), 7.44-7.37 (m, 3H), 6.99 (d, *J* = 9.0 Hz, 2H), 3.84-3.75 (m, 2H), 3.72-3.65 (m, 2H), 3.37-3.32 (m, 1H), 3.28-3.23 (m, 2H), 3.22-3.18 (m, 2H), 3.12-3.03 (m, 2H), 2.92 (t, *J* = 7.5 Hz, 2H), 1.84-1.77 (m, 2H), 1.70-1.63 (m, 2H), 1.46-1.40 (m, 8H), 1.24 (d, *J* = 7.5 Hz, 6H). $^{13}$C-NMR (126 MHz, MeOD): δ 158.08, 156.77, 149.97, 148.31, 138.58, 136.02, 132.89, 132.48, 127.67, 126.29, 126.05, 124.68, 118.51, 106.61, 58.01, 56.95, 53.10, 48.35, 40.69, 29.95, 28.54, 27.48, 27.32, 25.00, 15.44. HRMS (ESI+) m/z calculated for $C_{31}H_{45}ClN_7O_2S$+ $[M+H]^+$: 614.3038, found: 614.3050.

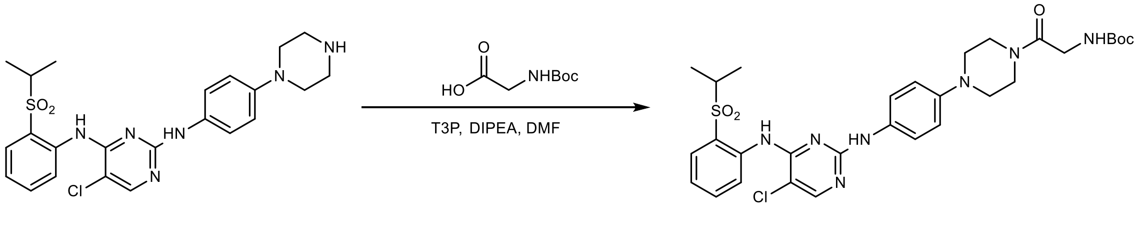

MKI-NH MKI-Gly-Boc

**tert-butyl (2-(4-(4-((5-chloro-4-((2-(isopropylsulfonyl)phenyl)amino)pyrimidin-2-yl)amino)phenyl)piperazin-1-yl)-2-oxoethyl)carbamate (MKI-Gly-Boc)**

Boc-Gly-OH (17.5 mg, 0.1 mmol, 1 equiv.), propanephosphonic acid anhydride (T3P) (108 mg, 0.17mmol, 1.7 equiv.) and diisopropylethylamine (52.3 µL, 0.3 mmol, 3 equiv.) were dissolved with dry DMF (3 mL) under $N_2$ followed by addition of MKI-NH (50 mg, 0.1 mmol, 1 equiv.) and stirred overnight. The reaction mixture was concentrated under reduced pressure and purified by preparative TLC (MeOH:DCM 1:9) to yield **MKI-Gly-Boc** as a yellow solid (36.6 mg, 57 µmol, 57%). $^1$H-NMR (500 MHz, $CDCl_3$): δ 9.63 (s, 1H), 8.58 (d, *J* = 8.5 Hz, 1H), 8.10 (s, 1H), 7.89 (dd, *J* = 8.0, 2.0 Hz, 1H), 7.55 (t, *J* = 8.5 Hz, 1H), 7.41 (d, *J* = 9.0 Hz, 2H), 7.27 (bs, 1H), 7.24 (t, *J* = 8.5 Hz, 2H), 6.88 (d, *J* = 9.0 Hz, 2H), 5.60 (s, 1H), 4.01 (d, *J* = 4.5 Hz, 2H), 3.81-3.79 (m, 2H), 3.57-3.55 (m, 2H), 3.25-3.20 (m, 1H), 3.14-3.10 (m, 4H), 1.45 (s, 9H), 1.30 (d, *J* = 6.5 Hz, 6H). $^{13}$C-NMR (126 MHz, $CDCl_3$): δ 167.06, 157.96, 155.99, 155.50, 154.98, 147.11, 138.50, 134.52, 132.76, 131.38, 124.57, 123.53, 123.26, 122.04, 117.67, 106.01, 79.87, 55.74, 50.36, 50.14, 44.50, 42.35, 42.07, 28.48, 15.46.

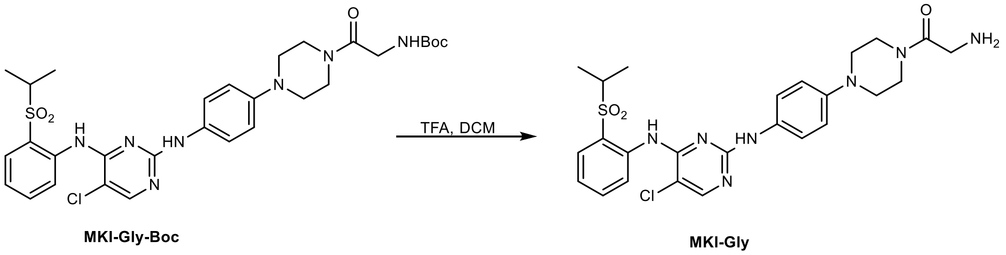


**2-amino-1-(4-(4-((5-chloro-4-((2-(isopropylsulfonyl)phenyl)amino)pyrimidin-2-yl)amino)phenyl)piperazin-1-yl)ethenone (MKI-Gly)**

MKI-Gly-Boc (60 mg, 0.093 mmol, 1 equiv.) was dissolved with DCM (1.5 mL) followed by a dropwise addition of TFA (375 µL). The mixture was stirred for 2h at room temperature and then concentrated under reduced pressure. The crude mixture was purified with reverse-phase chromatography on a Biotage C18 column (acetonitrile in $H_2O$, 5-95% gradient) to afford **MKI-Gly** as a yellow solid (16.7 mg, 30.6 µmol, 32%). $^1$H-

NMR (500 MHz, MeOD): δ 8.54 (d, *J* = 8.0 Hz, 1H), 8.11 (s, 1H), 7.90 (dd, *J* = 8.0, 2.0 Hz, 1H), 7.66 (dt, *J* = 9.0, 2.0 Hz, 1H), 7.40-7.35 (m, 3H), 6.98 (d, *J* = 8.5 Hz, 2H), 4.01 (s, 2H), 3.81-3.78 (m, 2H), 3.63-3.60 (m, 2H), 3.35-3.31 (m, 1H), 3.23-3.14 (m, 4H), 1.24 (d, *J* = 7.0 Hz, 6H). $^{13}$C-NMR (126 MHz, MeOD): δ 165.63, 163.40, 157.47, 152.49, 148.78, 139.08, 135.97, 133.40, 132.38, 126.84, 125.76, 125.34, 124.13, 118.59, 106.39, 56.93, 51.29, 51.02, 45.60, 43.16, 40.95, 15.46.

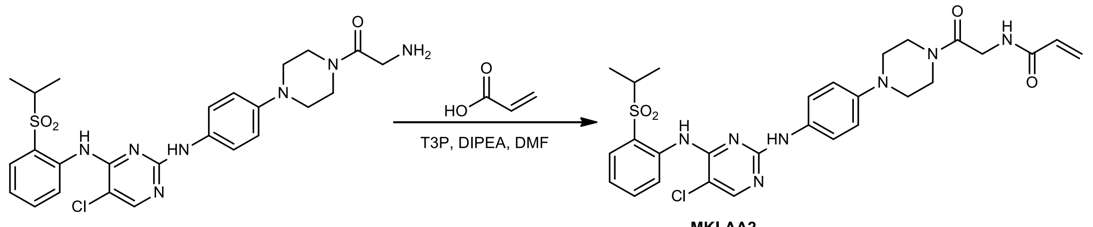


**N-(2-(4-(4-((5-chloro-4-((2-(isopropylsulfonyl)phenyl)amino)pyrimidin-2-yl)amino)phenyl)piperazin-1-yl)-2-oxoethyl)acrylamide (MKI-AA2)**

This compound was synthesized with the same procedure as MKI-AA, using acrylic acid (2.5 µL, 0.036 mmol, 1.2 equiv.), T3P (32 mg, 0.051 mmol, 1.7 equiv.), DIPEA (16 µL, 0.09 mmol, 3 equiv.), MKI-Gly (16.7 mg, 0.03 mmol, 1 equiv.) and DMF (2 mL). The preparative TLC was run with MeOH/DCM 1:9 to yield **MKI-AA2** as a yellow solid (6.0 mg, 1.67 µmol, 33%). $^{1}$H-NMR (500 MHz, $CDCl_3$): 9.65 (s, 1H), 8.58 (d, *J* = 8.5 Hz, 1H), 8.10 (s, 1H), 7.90 (dd, *J* = 8.0, 2.0 Hz, 1H), 7.56 (t, *J* = 8.5 Hz, 1H), 7.43 (d, *J* = 9.0 Hz, 2H), 7.25 (t, *J* = 8.0 Hz, 1H), 7.14 (bs, 1H), 6.90 (d, *J* = 8.5 Hz, 2H), 6.77 (bs, 1H), 6.32 (dd, *J* = 17.0, 1.5 Hz, 1H), 6.21 (dd, *J* = 17.0, 10.0 Hz, 1H), 5.69 (dd, *J* = 10.5, 1.5 Hz, 1H), 4.20 (d, *J* = 4.5 Hz, 2H), 3.84-3.81 (m, 2H), 3.63-3.60 (m, 2H), 3.28-3.19 (m, 1H), 3.17-3.12 (m, 4H), 3.31 (d, *J* = 7.0 Hz, 6H). $^{13}$C-NMR (126 MHz, $CDCl_3$): δ 166.51, 165.55, 157.89, 155.57, 154.88, 147.10, 138.51, 134.53, 132.80, 131.43, 130.50, 127.05, 124.67, 123.56, 123.33, 122.04, 117.78, 106.17, 55.78, 50.43, 50.18, 44.60, 42.22, 41.42, 15.49. HRMS (ESI+) m/z calculated for $C_{28}H_{32}ClN_7O_4S$ $[M+H]^+$: 598.19978; found: 598.19870.

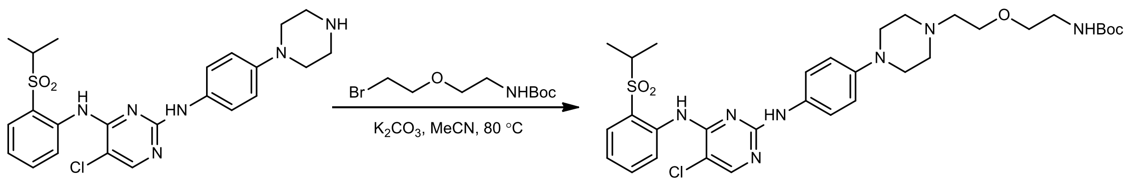


**tert-butyl (2-(2-(4-(4-((5-chloro-4-((2-(isopropylsulfonyl)phenyl)amino)pyrimidin-2-yl)amino)phenyl)piperazin-1-yl)ethoxy)ethyl)carbamate (MKI-PEG1-Boc)**

MKI-NH (100 mg, 0.17 mmol, 1 equiv.), N-Boc-PEG1-bromide (66 mg, 0.2 mmol, 1.2 equiv.) and potassium carbonate (83 mg, 0.5 mmol, 3 equiv.) were dissolved with 2.5 acetonitrile and refluxed at 80 °C for overnight. The mixture was diluted with dichloromethane and filtered. The filtrate was concentrated under reduced pressure and purified by preparative TLC (MeOH:DCM 1:9) to yield **MKI-PEG1-Boc** as a yellow solid (82.7 mg, 122 µmol, 61%). $^1$H-NMR (500 MHz, $CDCl_3$): δ 9.60 (s, 1H), 8.59 (d, *J* = 8.5 Hz, 1H), 8.09 (s, 1H), 7.87 (d, *J* = 8.0 Hz, 1H), 7.53 (t, *J* = 7.5 Hz, 1H) 7.36 (d, *J* = 8.5 Hz, 2H), 7.20 (t, *J* = 7.5 Hz, 2H), 6.88 (d, *J* = 8.5 Hz, 2H), 5.23 (bs, 1H), 3.62 (t, *J* = 6.0 Hz, 2H), 3.53 (t, *J* = 5.0, 2H), 3.33-3.29 (m, 2H), 3.25-3.20 (m, 1H), 3.20-3.18 (m, 4H), 2.69-2.67 (m, 4H), 2.64 (t, *J* = 5.5 Hz, 2H), 1.42 (s, 9H), 1.29 (d, *J* = 7.0 Hz, 6H). $^{13}$C-NMR (126 MHz, $CDCl_3$): δ 158.34, 156.13, 155.43, 155.32, 147.84, 138.62, 134.56, 131.68, 131.28, 124.35, 123.50, 123.05, 122.34, 116.73, 105.73, 79.32, 69.97, 68.22, 57.82, 55.68, 53.66, 49.72, 40.46, 28.56, 15.45.

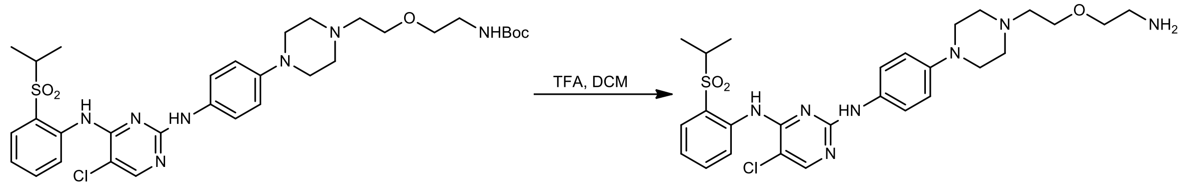


**N2-(4-(4-(2-(2-aminoethoxy)ethyl)piperazin-1-yl)phenyl)-5-chloro-N-4-(2-(isopropylsulfonyl)phenyl)pyrimidine-2,4-diamine (MKI-PEG1)**

MKI-PEG1-Boc (82.7 mg, 0.012 mmol, 1 equiv.) was dissolved with DCM (2 mL) followed by a dropwise addition of TFA (492 µL). The mixture was stirred for 2h at room temperature and then concentrated under reduced pressure. The crude mixture was purified with reverse-phase chromatography on a Biotage C18 column (acetonitrile in $H_2O$, 5-95% gradient) to afford **MKI-PEG1** in quantitative yield as a yellow solid. $^1$H-NMR (500 MHz, MeOD): δ 8.43 (d, *J* = 8.5 Hz, 1H), 8.14 (s, 1H), 7.91 (dd, *J* = 7.5, 1.5 Hz, 1H), 7.66 (t, *J* = 7.0 Hz, 1H), 7.42 (dt, *J* = 7.5, 1.0 Hz, 1H), 7.37 (d, *J* = 8.5 Hz, 2H), 6.98 (d, *J* = 9.0 Hz, 2H), 3.93 (t, *J* = 4.5 Hz, 2H), 3.78 (t, *J* = 5.0 Hz, 2H), 3.73 (bs, 2H), 3.49 (t, *J* = 5.0 Hz, 2H), 3.38-3.32 (m, 1H), 3.19 (t, *J* = 5.0 Hz, 2H), 1.23 (d, *J* = 7.0 Hz, 6H). $^{13}$C-NMR (126 MHz, MeOD): δ 158.84, 154.53, 149.17, 145.49, 137.63, 136.09, 132.56, 131.22, 128.54, 127.07, 126.85, 125.49, 118.32, 106.76, 68.01, 65.36, 56.98, 54.75, 53.27, 47.72, 40.27, 15.38.

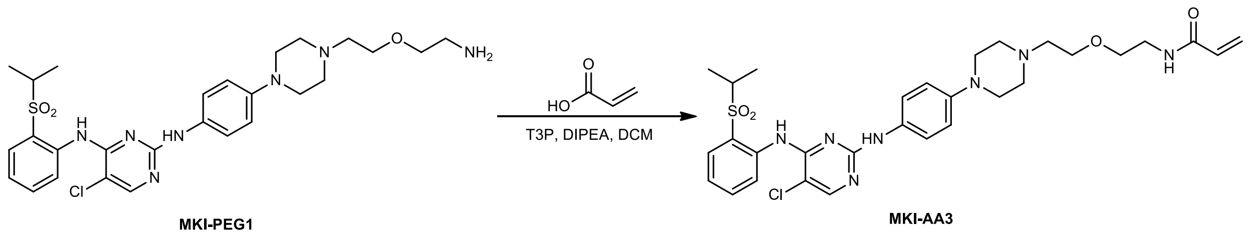


**N-(2-(2-(4-(4-((5-chloro-4-((2-(isopropylsulfonyl)phenyl)amino)pyrimidin-2-yl)amino)phenyl)piperazin-1-yl)ethoxy)ethyl)acrylamide (MKI-AA3)**

This compound was synthesized with the same procedure as MKI-AA, using acrylic acid (4.3 µL, 0.068 mmol, 1.2 equiv.), T3P (61 mg, 0.097 mmol, 1.7 equiv.), DIPEA (29 µL, 0.17 mmol, 3 equiv.), MKI-PEG1 (33 mg, 0.057 mmol, 1 equiv.) and DMF (2 mL). The preparative TLC was run with MeOH/DCM 5:95 to yield **MKI-AA3** as a yellow solid (3.7 mg, 5.9 µmol, 10%). $^1$H-NMR (500 MHz, $CDCl_3$): δ 9.61 (s, 1H), 8.59 (d, *J* = 8.5 Hz, 1H), 8.10 (s, 1H), 7.89 (dd, *J* = 7.5, 1.5 Hz, 1H), 7.56 (dt, *J* = 8.0, 1.5 Hz, 1H), 7.49 (d, *J* = 9.0 Hz, 2H), 7.23 (dt, *J* = 7.5, 1.5 Hz, 1H), 6.95 (bs, 1H), 6.89 (d, *J* = 8.5 Hz, 2H), 6.30 (dd, *J* = 17.0, 1.5 Hz, 1H),6.18-6.12 (m, 1H), 5.62 (dd, *J* = 10.0, 1.5 Hz, 1H), 3.69-3.66 (m, 2H), 3.61-3.59 (m, 2H), 3.57-3.53 (m, 2H), 3.26-3.20 (m, 4H), 2.81-2.71 (m, 6H), 1.31 (d, *J* =

7.0 Hz, 6H). $^{13}C$-NMR (126 MHz, $CDCl_3$): δ 168.57, 165.73, 158.23, 155.52, 155.32, 138.63, 134.58, 132.02, 131.38, 131.06, 126.53, 124.54, 123.55, 123.18, 122.23, 117.04, 106.01, 69.69, 57.82, 55.74, 53.67, 49.53, 39.26, 29.85, 15.51. HRMS (ESI+) m/z calculated for $C_{30}H_{38}ClN_7O_4S$ $[M+H]^+$: 628.24673; found: 628.24895.

### $^{1}$H-NMR for MKI-AA (500 MHz, $CDCl_3$)

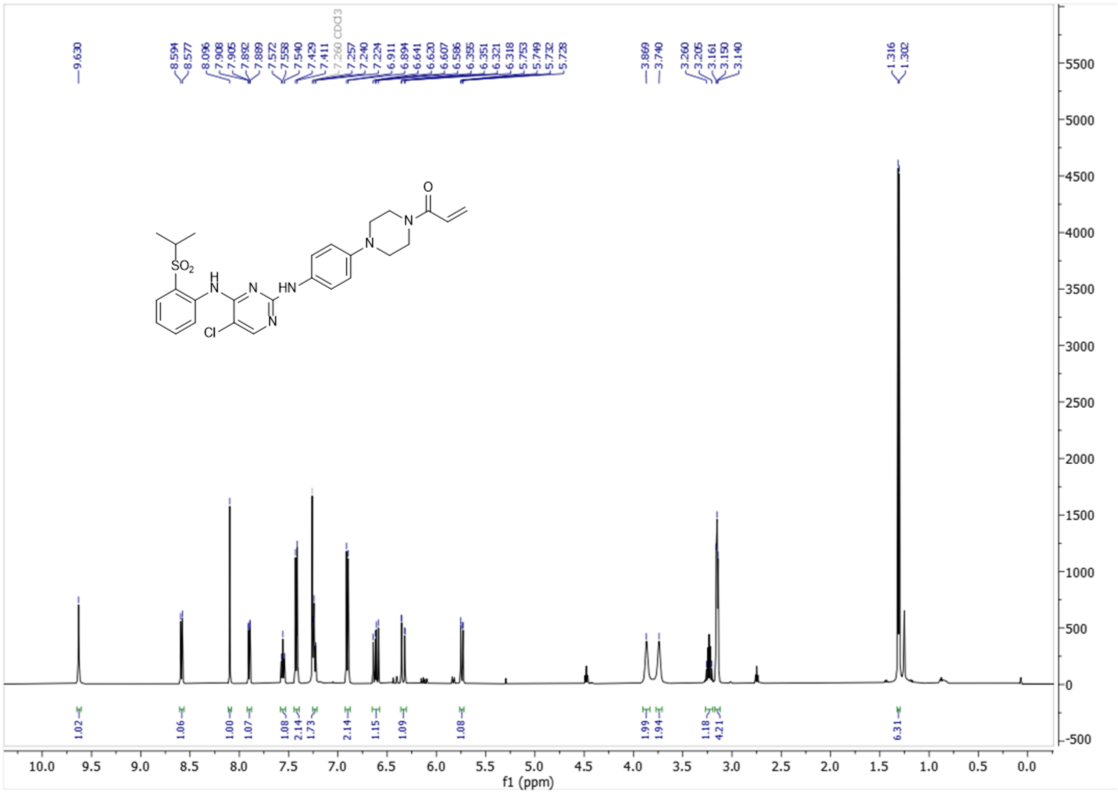


### $^{13}$C-NMR for MKI-AA (126 MHz, $CDCl_3$)

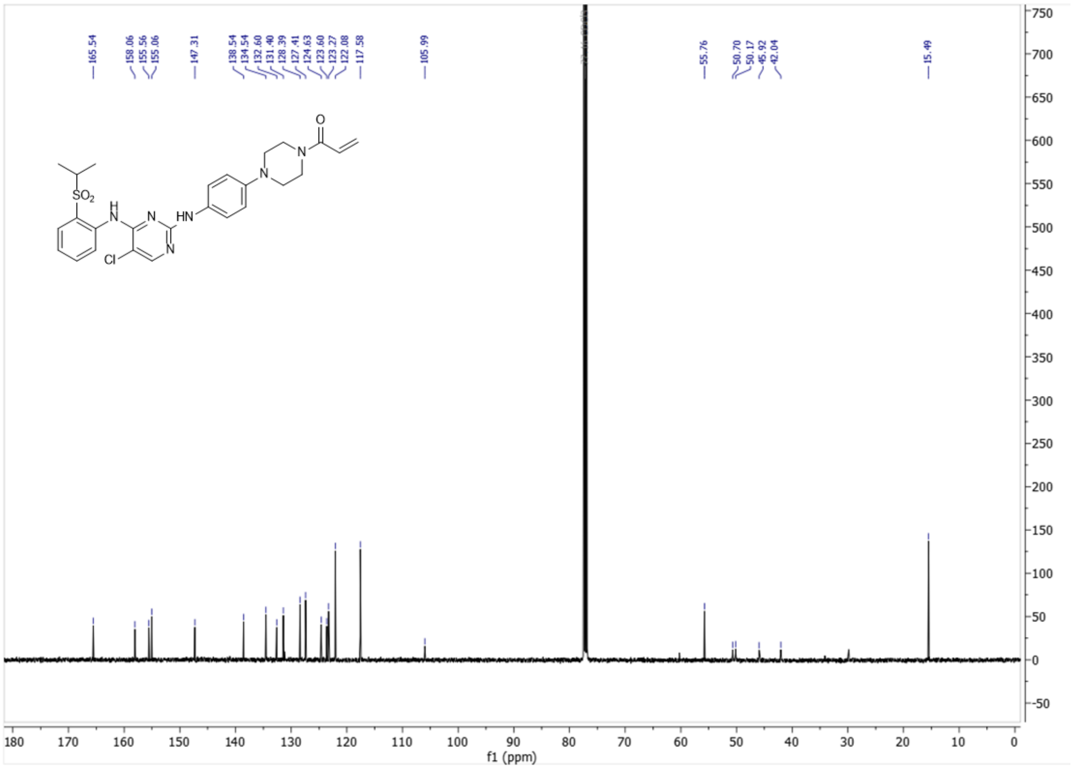

**$^1$H-NMR for MKI-CA (500 MHz, $CDCl_3$)**

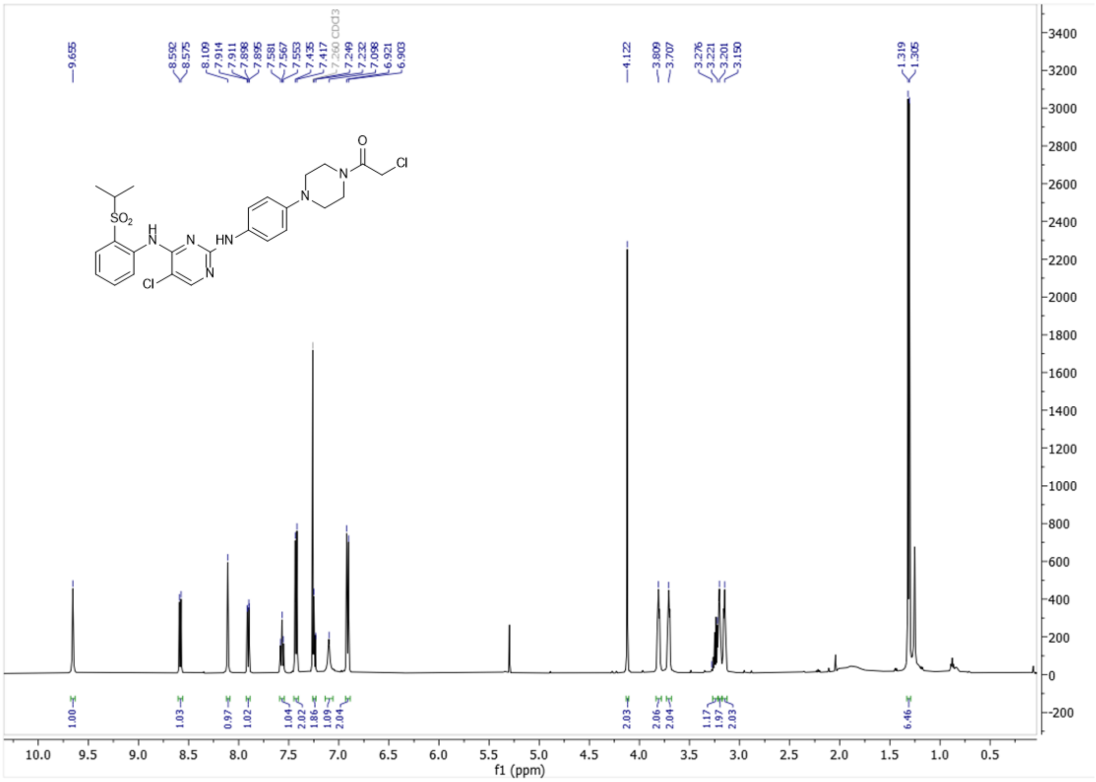


**$^{13}$C-NMR for MKI-CA (126 MHz, $CDCl_3$)**

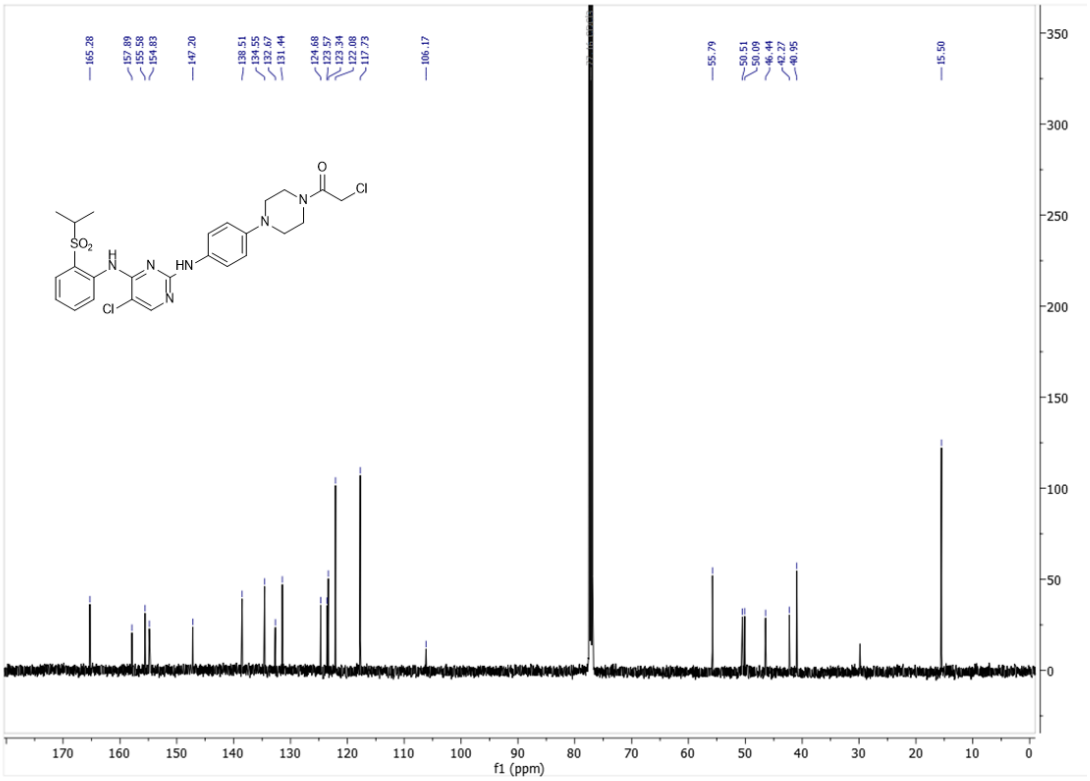

### $^1$H-NMR for MKI-PA (500 MHz, $CDCl_3$)

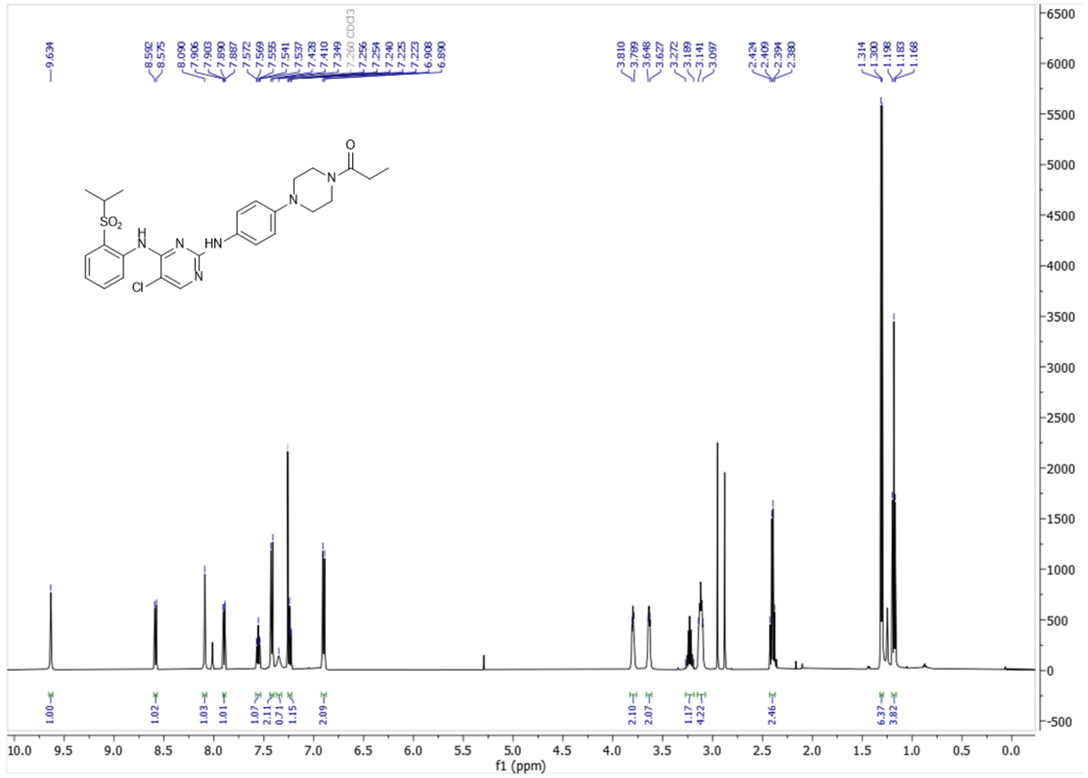

### $^{13}$C-NMR for MKI-PA (126 MHz, $CDCl_3$)

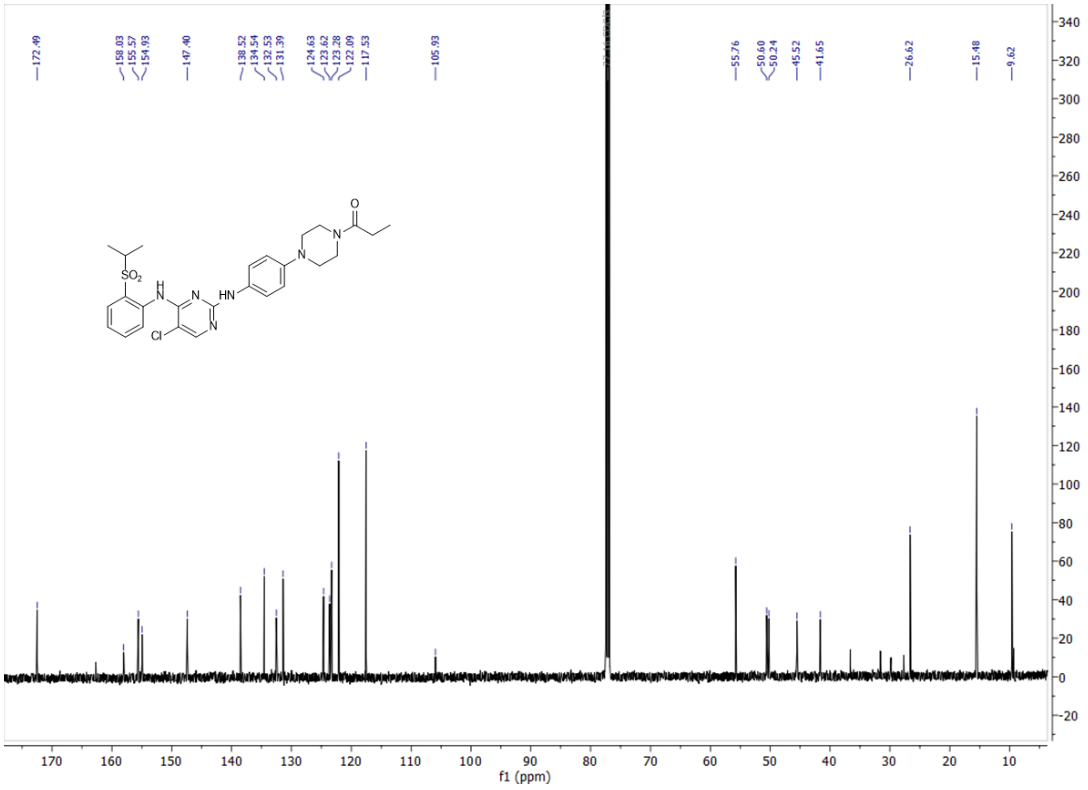

### $^{1}$H-NMR for MKI-A-Boc (500 MHz, $CDCl_3$)

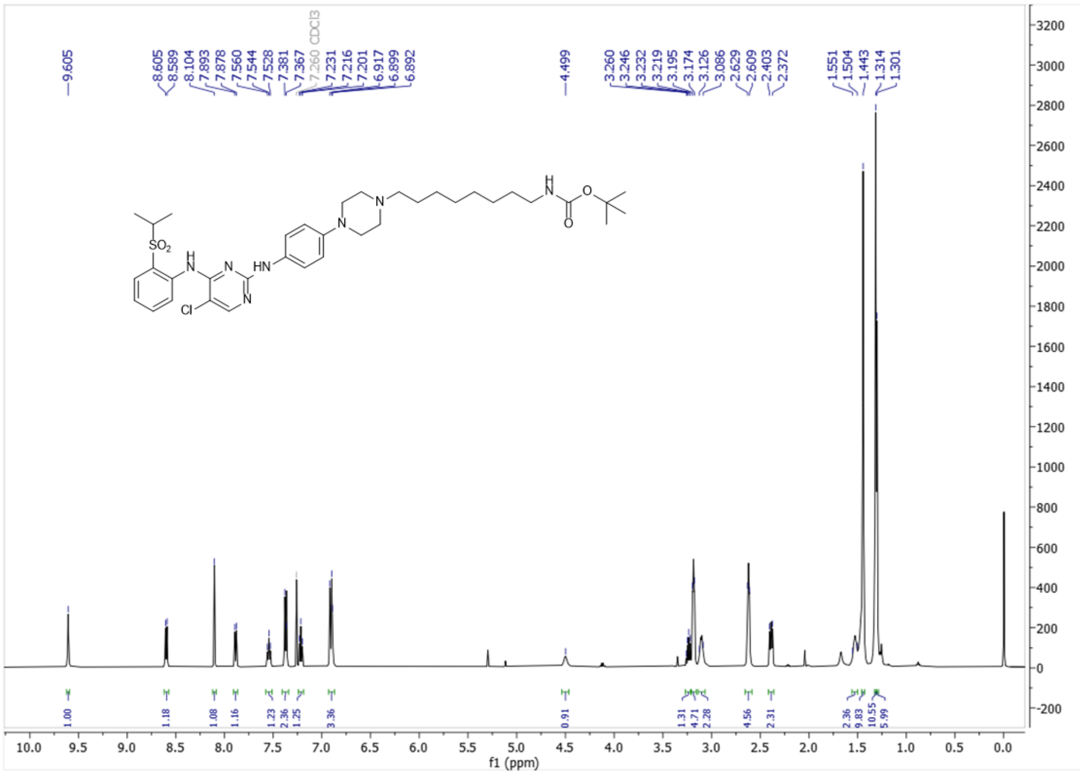

### $^{13}$C-NMR for MKI-A-Boc (126 MHz, $CDCl_3$)

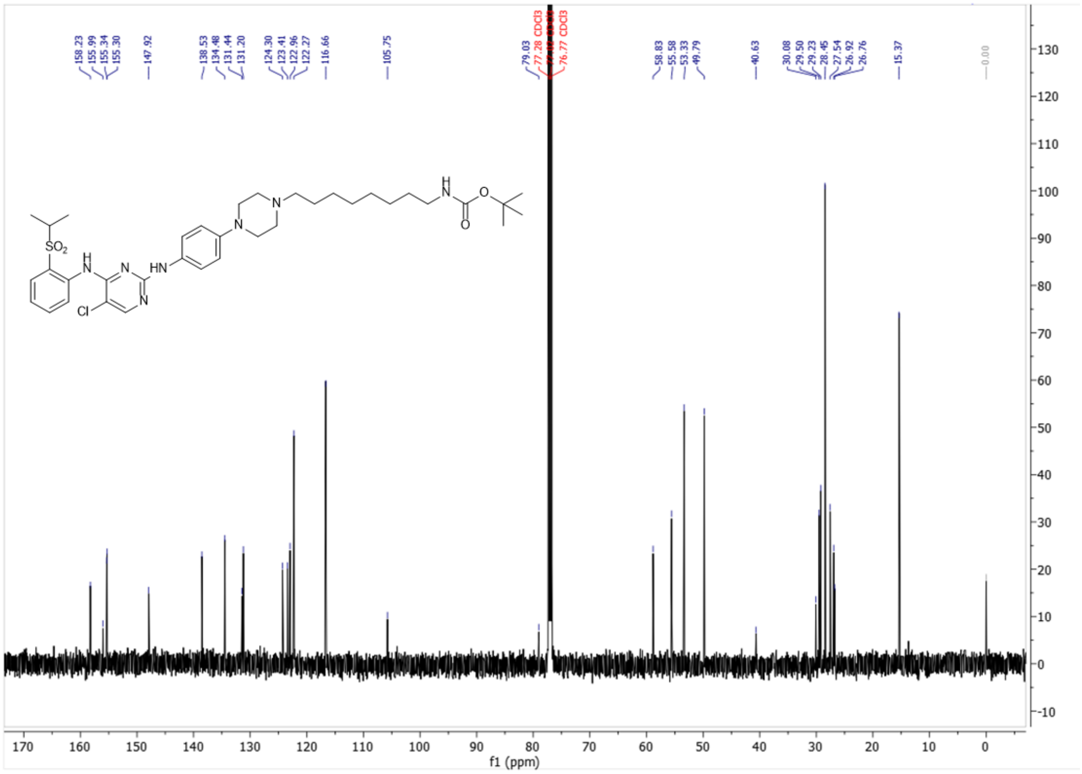

### $^{1}$H-NMR for MKI-A (500 MHz, MeOD)

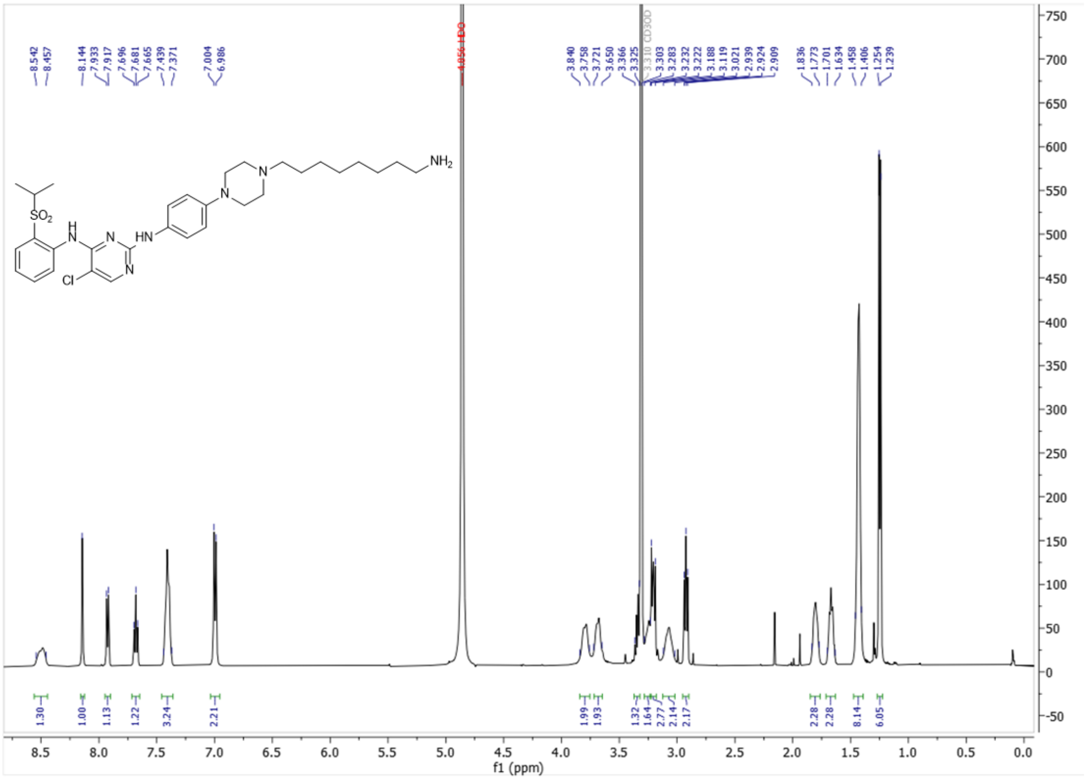


### $^{13}$C-NMR for MKI-A (126 MHz, MeOD)

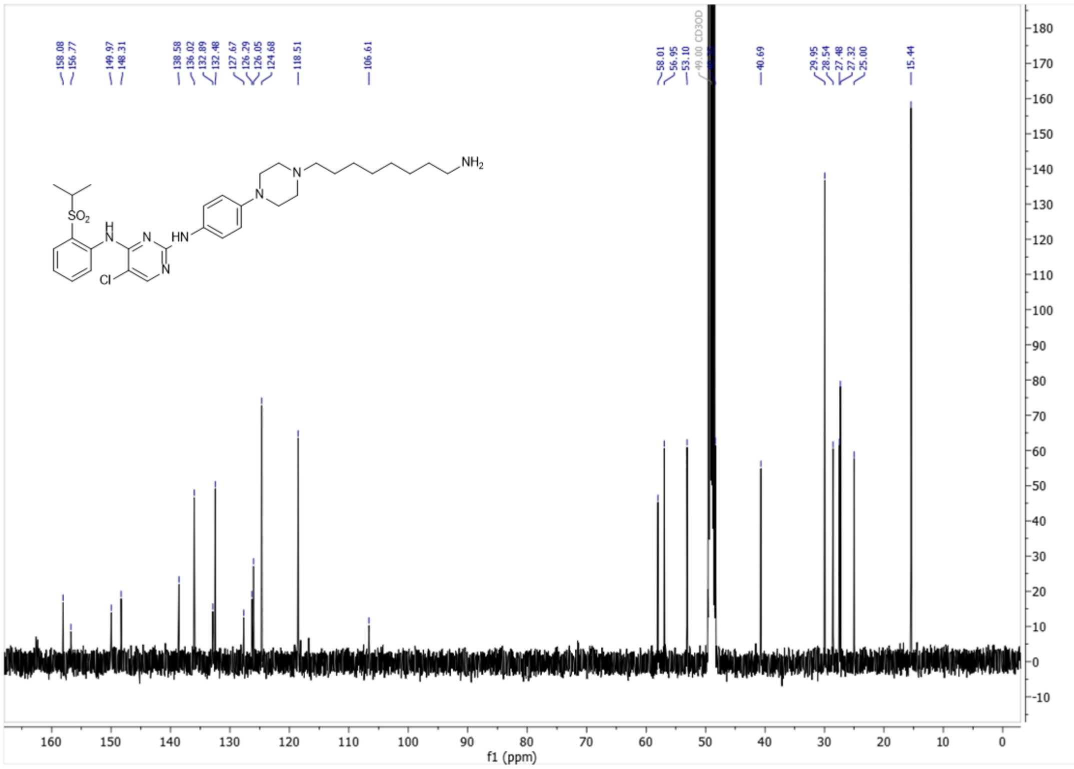

### $^{1}$H-NMR for MKI-Gly-Boc (500 MHz, $CDCl_3$)

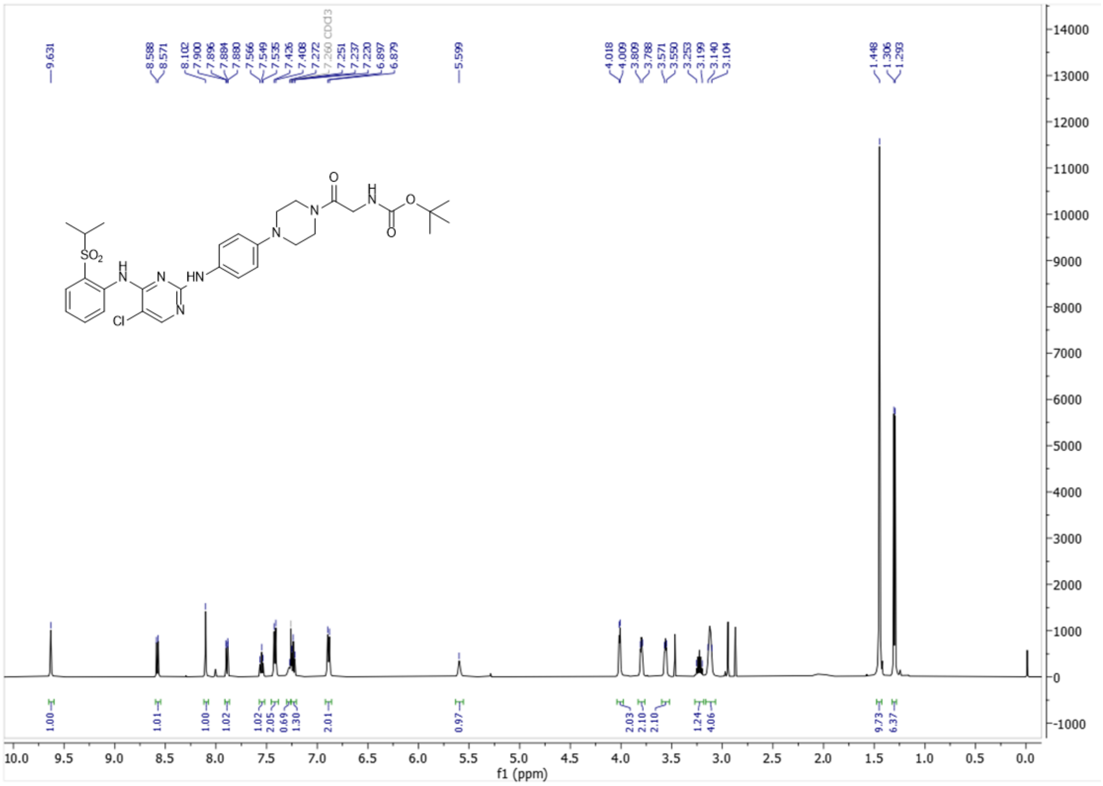


### $^{13}$C-NMR for MKI-Gly-Boc (126 MHz, $CDCl_3$)

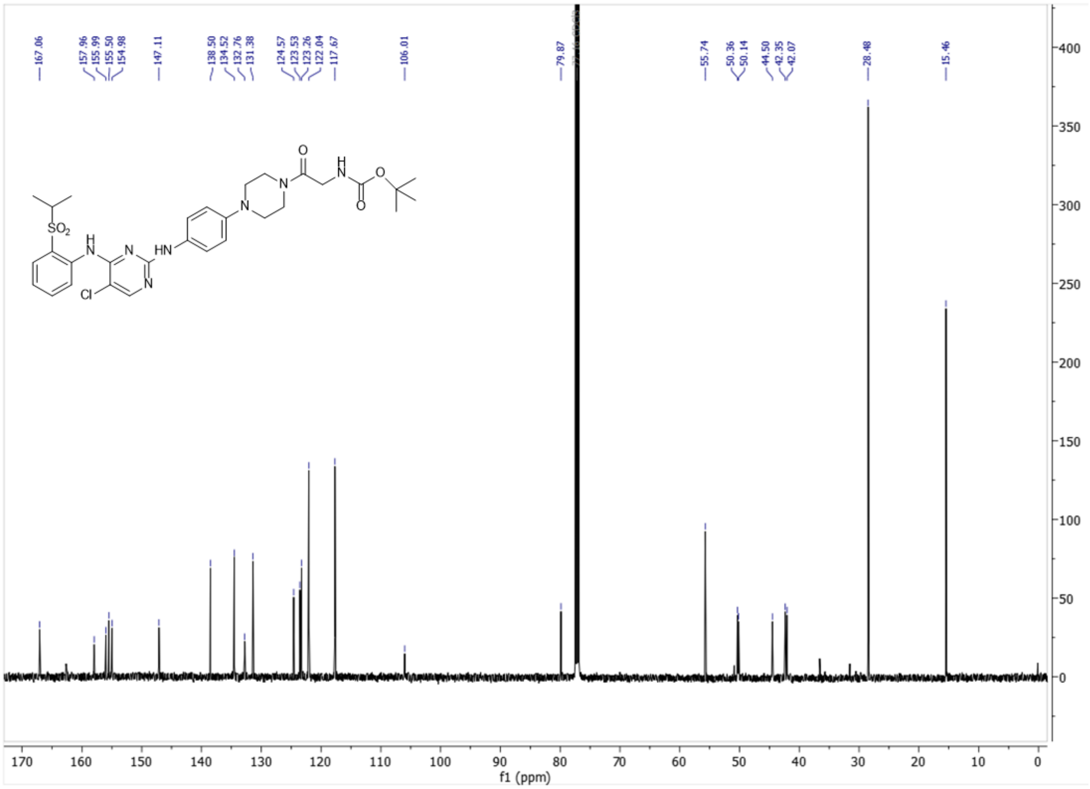

**$^{1}$H-NMR for MKI-Gly (500 MHz, MeOD)**

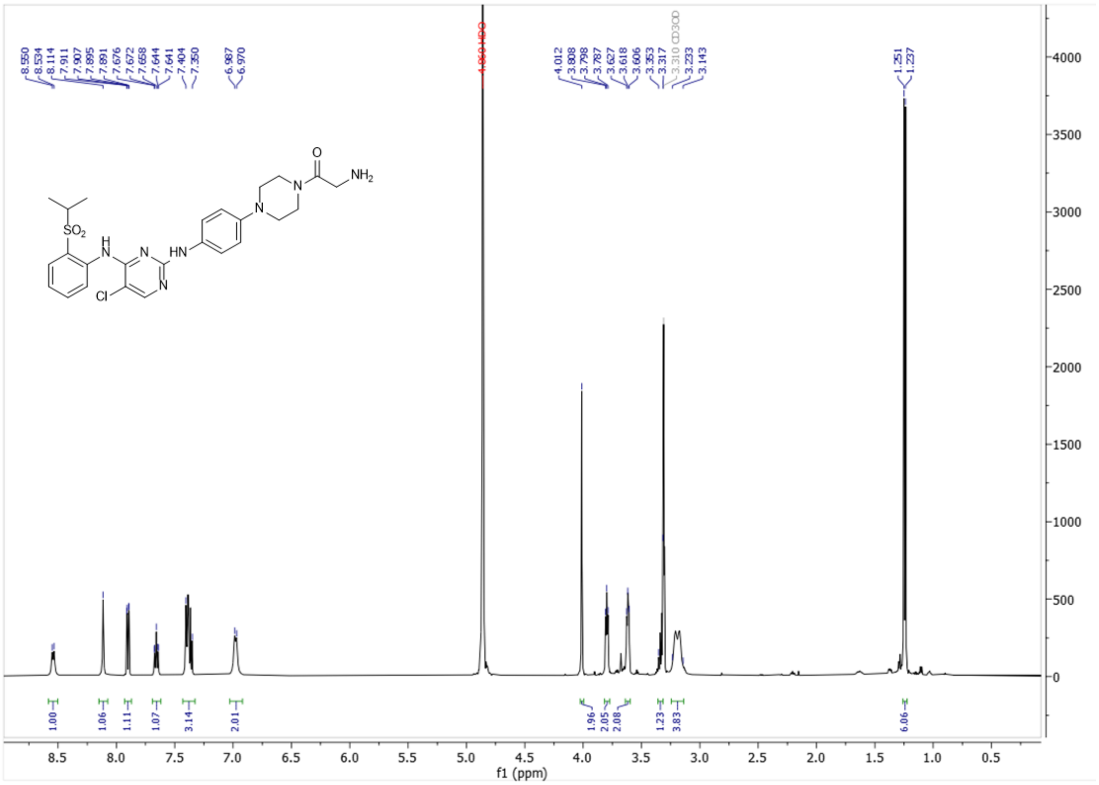


**$^{13}$C-NMR for MKI-Gly (126 MHz, MeOD)**

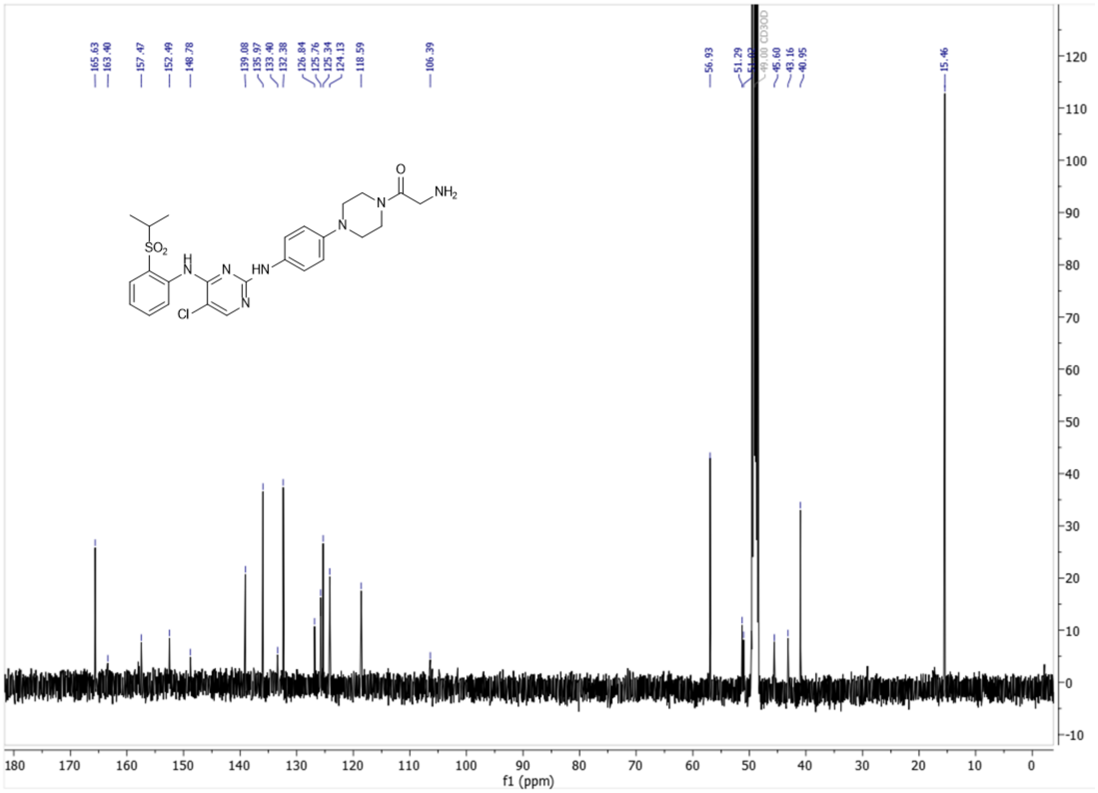

#### $^{1}$H-NMR for MKI-AA2 (500 MHz, $CDCl_3$)

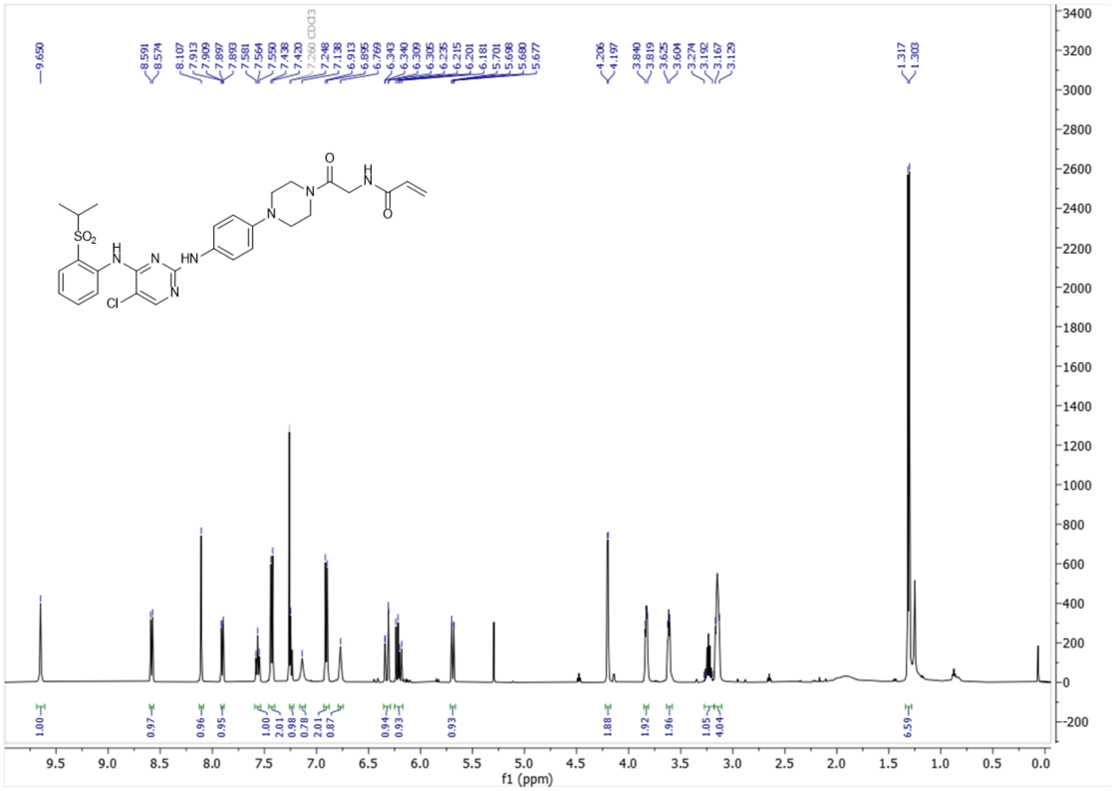


#### $^{13}$C-NMR for MKI-AA2 (126 MHz, $CDCl_3$)

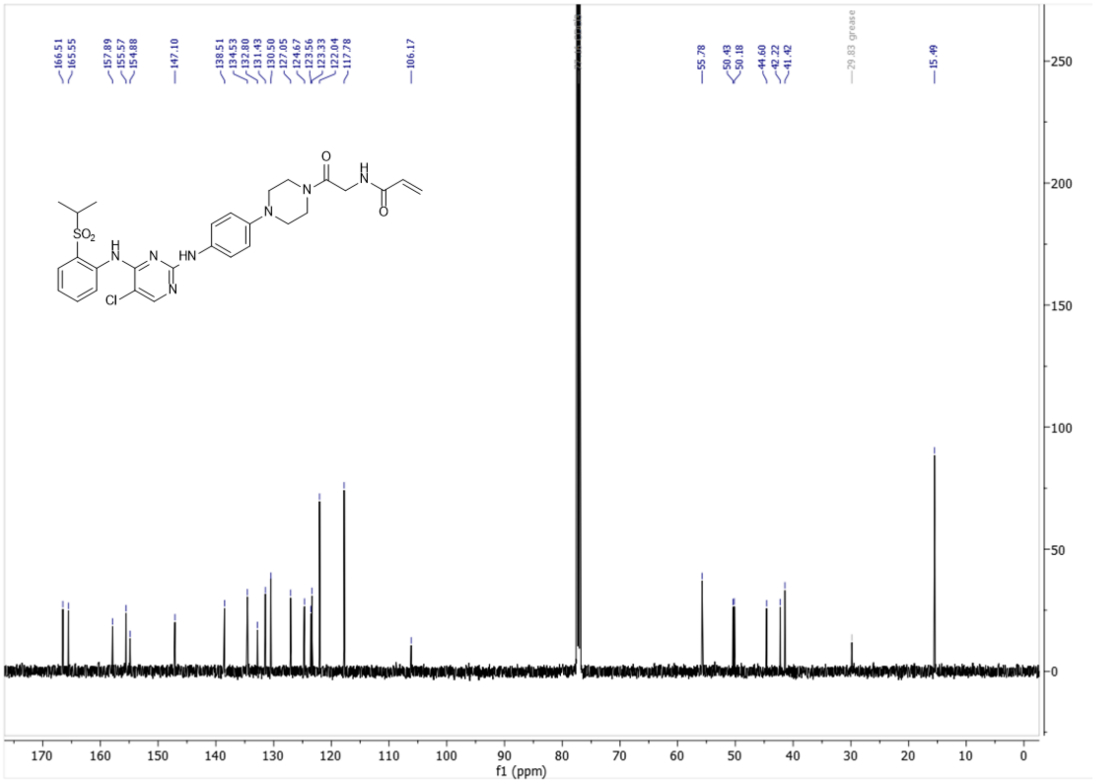

$^{1}$H-NMR for MKI-PEG1-Boc (500 MHz, CDCl$_3$)

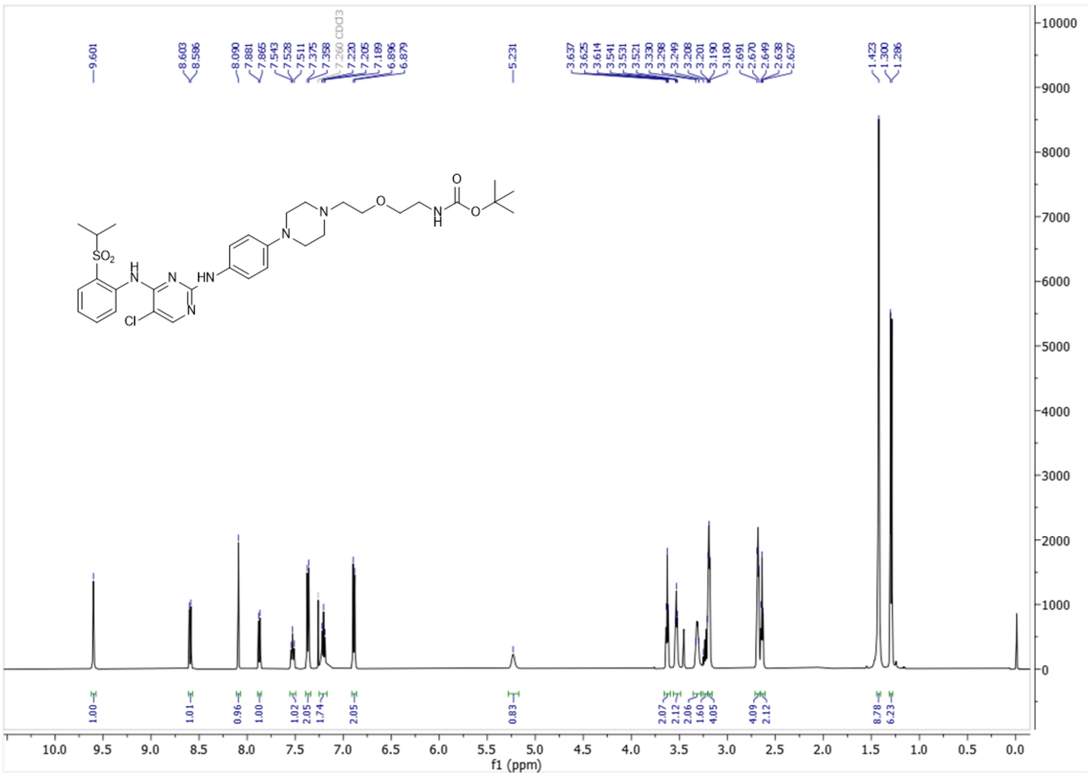

$^{13}$C-NMR for MKI-PEG1-Boc (126 MHz, CDCl$_3$)

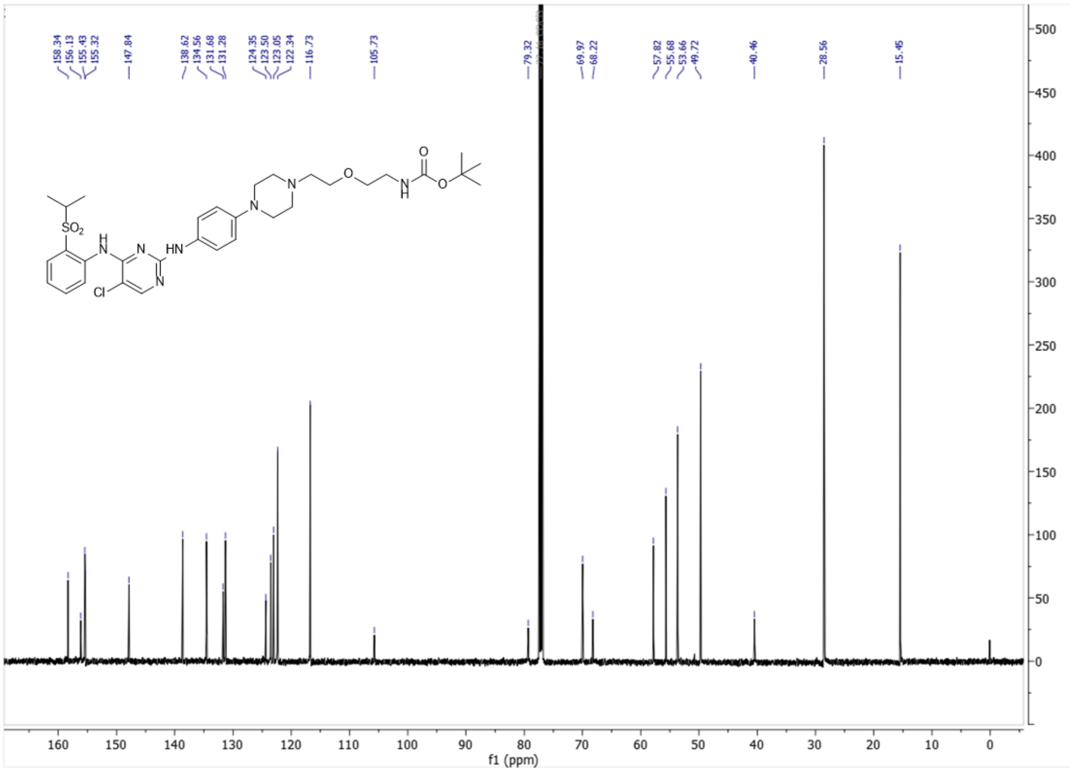

### [1]H-NMR for MKI-PEG1 (500 MHz, MeOD)

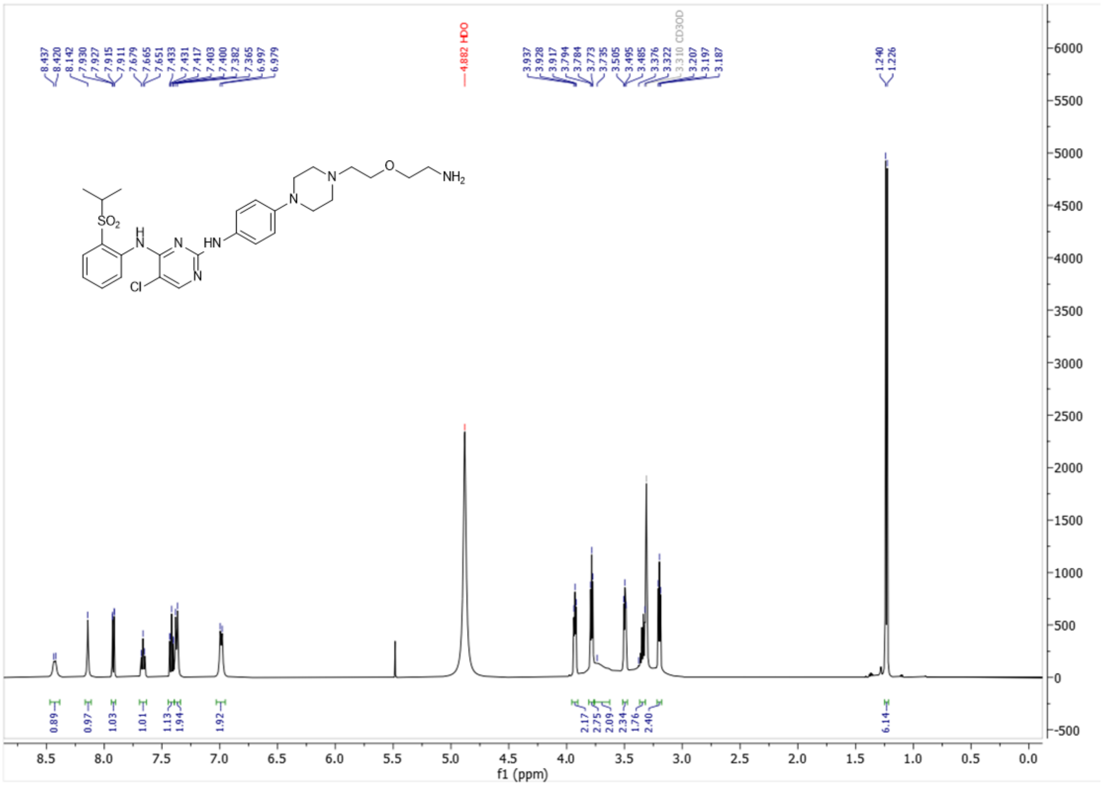


### [13]C-NMR for MKI-PEG1 (126 MHz, MeOD)

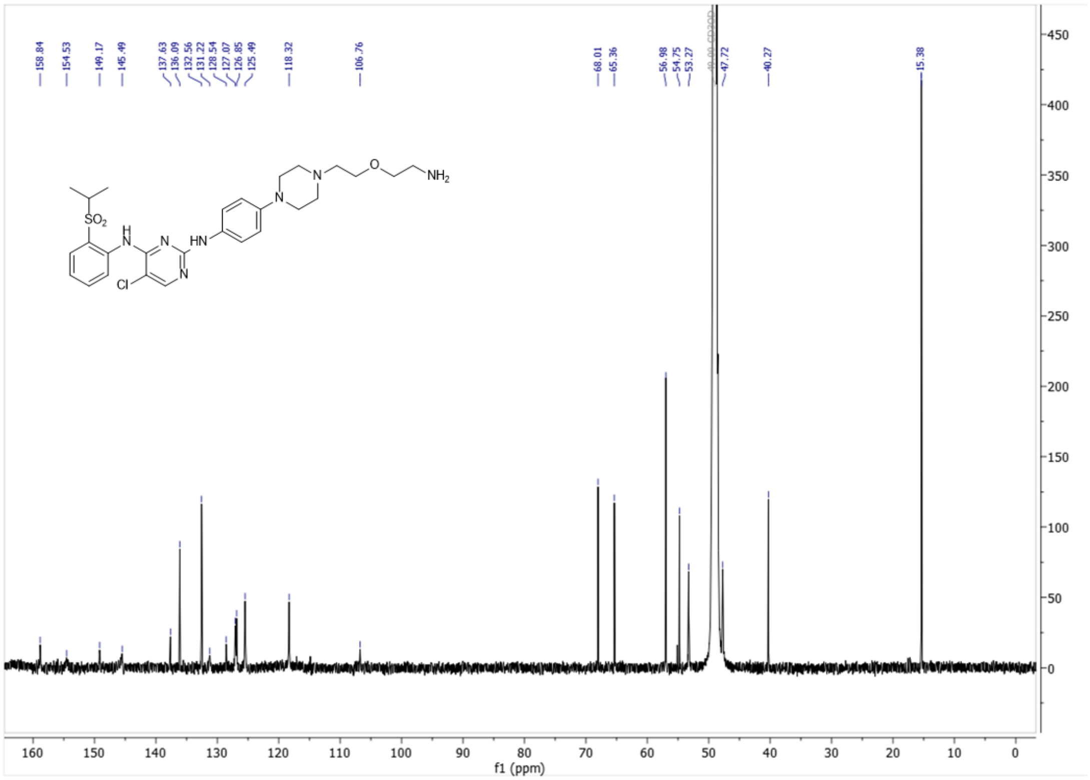

**$^1$H-NMR for MKI-AA3 (500 MHz, $CDCl_3$)**

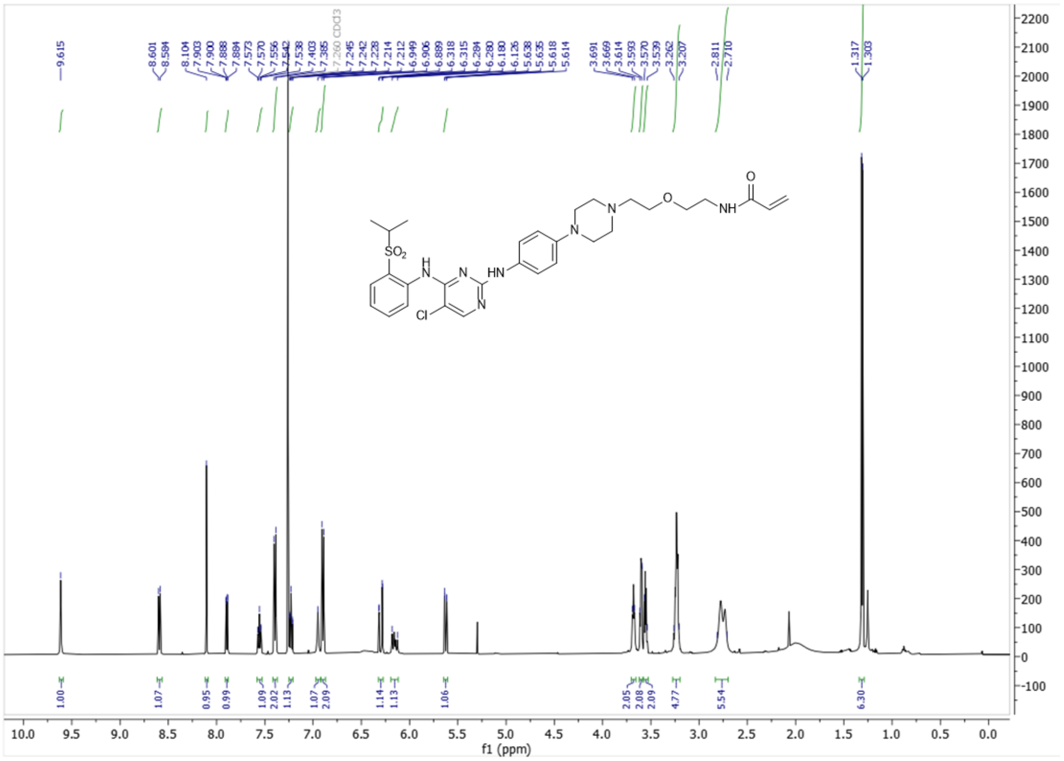

**$^{13}$C-NMR for MKI-AA3 (126 MHz, $CDCl_3$)**

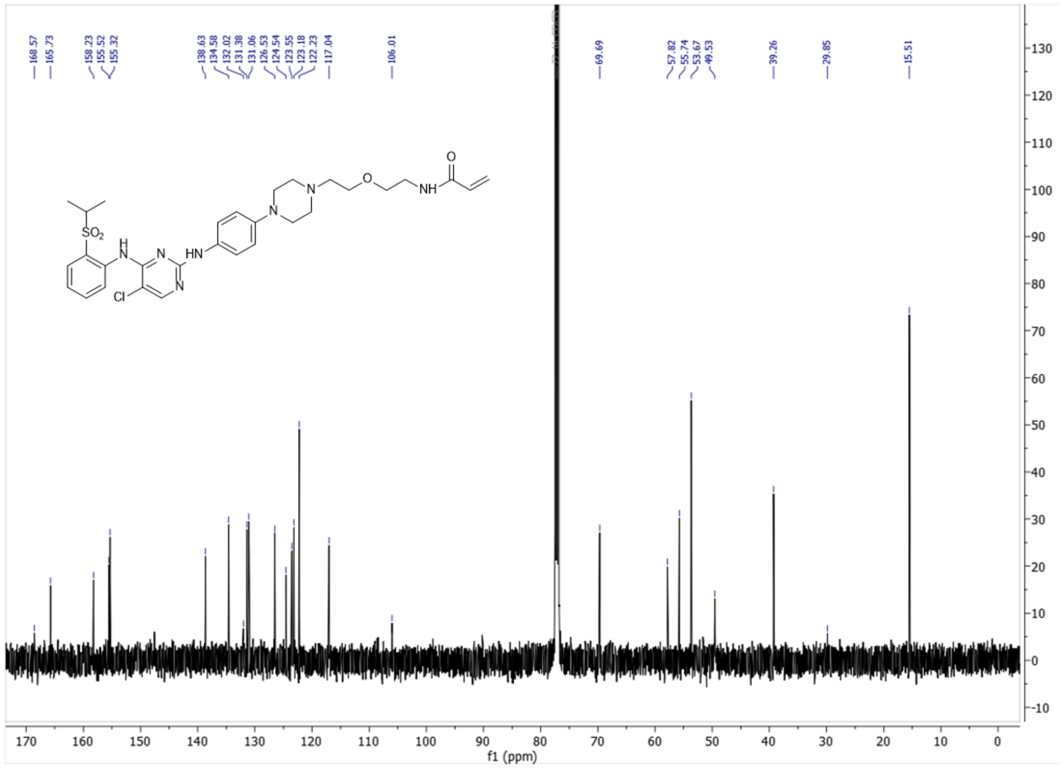
